# Endocardium-to-coronary artery differentiation during heart development and regeneration involves sequential roles of Bmp2 and Cxcl12/Cxcr4

**DOI:** 10.1101/2021.10.25.465773

**Authors:** Gaetano D’Amato, Ragini Phansalkar, Jeffrey A. Naftaly, Pamela E. Rios Coronado, Dale O. Cowley, Kelsey E. Quinn, Bikram Sharma, Kathleen M. Caron, Alessandra Vigilante, Kristy Red-Horse

## Abstract

Regenerating coronary blood vessels has the potential to ameliorate ischemic heart disease, yet there is currently no method of stimulating clinically effective cardiac angiogenesisis. Endocardial cells— a particularly plastic cell type during development—line the heart lumen and are natural coronary vessel progenitors. Their intrinsic angiogenic potential is lost in adults, but studying the endocardial-to-coronary developmental pathway could identify methods of re-instating this process in diseased hearts. Here, we use a combination of mouse genetics and scRNAseq of lineage-traced endothelial cells to identify novel regulators of endocardial angiogenesis and precisely assess the role of Cxcl12/Cxcr4 signaling. Time-specific lineage tracing demonstrated that endocardial cells differentiated earlier than previously thought, largely at mid-gestation. A new mouse line reporting the activity of Cxcr4 revealed that, despite widespread Cxcl12 and Cxcr4 expression, only a small subset of these coronary endothelial cells activated the receptor, which were mostly in arteries. In accordance with these two findings, *Cxcr4* deletion in the endocardial lineage only affected artery formation and only when deleted before mid-gestation. Integrating scRNAseq data of coronary endothelial cells from the endocardial lineage at both mid- and late-gestation identified a transitioning population that was specific to the earlier timepoint that specifically expressed Bmp2. Recombinant Bmp2 stimulated endocardial angiogenesis in an in vitro explant assay and in neonatal mouse hearts upon myocardial infarction. Our data shed light on how understanding the molecular mechanisms underlying endocardial-to-coronary transitions can identify new potential therapeutic targets that could promote revascularization of the injured heart.

## INTRODUCTION

Ischemic heart disease—the leading cause of death worldwide—most commonly results from occlusion of the coronary arteries and can lead to myocardial infarction (MI) and heart failure (Dunbar et al. 2018). One intriguing therapeutic approach to ameliorate heart function upon MI could be to rebuild damaged coronary blood vessels. However, developing a medical intervention that regenerates coronary vasculature has proven difficult (Ylä-Herttuala et al. 2017), likely due to our poor understanding of how cardiac-specific blood vessels develop and respond to injury.

During development, the majority of coronary vessels endothelial cells (CV ECs) arise from two spatially unrelated cell types: endothelial cells (ECs) from the sinus venosus (SV), the venous inflow tract of the developing heart, and endocardium (endo), the innermost layer of embryonic and adult heart (Red-Horse et al. 2010; Wu et al. 2012; Tian et al. 2013; Chen et al. 2014b; Tian et al. 2014; Zhang et al. 2016; Sharma et al. 2017c; Su et al. 2018b). There is a consensus that the initial sprouting angiogenesis from SV or endo contributes to complementary regions of the vascular plexus (Tian et al. 2013; Chen et al. 2014a; Tian et al. 2014; Sharma et al. 2017b) and is driven by different signaling pathways, i.e. epicardial VEGF-C/ELA versus intramyocardial VEGFA, respectively (Wu et al. 2012; Chen et al. 2014b; Sharma et al. 2017c; Neal et al. 2019). The presence of dual, unrelated progenitors for CV ECs within the developing heart may have several biological benefits. It ensures fast vascularization of the growing heart during mid-gestation (Rhee et al. 2018) and provides alternative progenitors when angiogenesis from one source is abrogated (Sharma et al. 2017c).

Over the last ten years, mouse genetics, lineage tracing analysis, and single-cell RNA sequencing (scRNAseq) have been used to extensively study how CV ECs sprout and migrate from the SV to form an immature vascular plexus that eventually differentiates into mature capillary, veins, and arteries (Sharma et al. 2017a; Su et al. 2018a; Zhang et al. 2018). By comparison, there has been much less focus on endo sprouting, and there are major gaps in our knowledge on this process.

Although molecular details are sparse, sequential lineage tracing efforts over the last decade have progressively refined our understanding of when and where in the heart endo cells contribute to CV ECs in mice. Initially, *Nfatc1-Cre* lineage labeling suggested the endo to be the major source of intramyocardial capillaries and coronary arteries, while concomitant EC clonal analysis indicated that most of this contribution occurred in and near the septum (Red-Horse et al. 2010; Wu et al. 2012). Subsequently, new *Cre* lines allowing more specific spatiotemporal lineage labeling reported that the endo minimally contributed to CV ECs within ventricular free walls before birth, producing mostly septal CV ECs at this stage (Zhang et al. 2016). These same tools also showed that a second wave of endo-derived CV ECs begin to populate the inner half of the free walls shortly after birth (Tian et al. 2014). From histological analyses, the authors concluded that this wave of CV ECs arose from endo cells that become trapped during the process of neonatal trabecular compaction, although alternative interpretations existed since endo in these experiments were marked at embryonic day (E)8. Indeed, a recent study employing a lineage labeling time course revealed that the second wave derives from endo cells that differentiated into CV ECs at some point prior to E16.5 (Lu et al. 2021).

The restriction of endo differentiation to embryonic stages is consistent with studies showing that angiogenesis in response to cardiac injury does not normally occur from the endo in neonate and adult hearts (Tang et al. 2018; Lu et al. 2021). However, some can be induced by Vegf-B or Vegfr2 overexpression after experimental myocardial infarction (MI) (Jiang et al. 2021; Räsänen et al. 2021). Re-activating endo angiogenesis in adults could be an attractive therapeutic strategy, particularly since ischemic injuries are frequently localized to subendocardial myocardium. To reach this goal we must learn more about normal development. Current critical outstanding questions include: at which precise time points between E8 and 16.5 do endo cells differentiate into CV ECs and what are the molecular players.

Here, we used an expanded lineage tracing time-course strategy and multi-stage scRNAseq to further define the endo angiogenic window and discover molecular regulators. We show that the endo-to-CV ECs transition largely occurs between E10.5-13.5. Endo-derived CVs first produce septal CV ECs that subsequently migrate outward to form vessel plexuses outside the septum. Ligand-receptor expression patterns suggested that this migration involved Cxcl12/Cxcr4, but mouse genetics and a new mouse line that reports active Cxcr4 signaling showed that this axis functions primarily during artery formation. To look for regulators of early steps in endo angiogenesis, we contrasted scRNAseq datasets containing labeled endo-derived CV ECs from early and late embryonic development. *Bmp2* was upregulated in a subset of CV ECs transitioning from the endo, which was present in early but not late developmental time points. Recombinant Bmp2 potentiated Vegf-A induced angiogenesis, and forced *Bmp2* expression in neonatal hearts stimulated endo angiogenesis after MI. Collectively, our data highlight how spatiotemporally restricted signaling influences the capability of endo cells to differentiate into CVs and that Bmp2 can re-activate this activity within favorable angiogenic/hypoxic environments. These findings could inspire progenitor cell-based therapy for neovascularization following ischemic heart disease.

## RESULTS

### Differentiation of endo to CV ECs is largely restricted to early heart development

Endo cells are known to produce CV ECs that first appear during mid-gestation embryogenesis through angiogenic sprouting and continue to increase in number during postnatal life. It was posited that differentiation of endo cells into CV ECs occurs throughout this entire time period due to cell capture during trabecular compaction (Tian, 2014). However, recent studies show that the actual cell fate conversion from endo into CV ECs is restricted to embryonic stages (Lu et al. 2021). One unanswered question is at what stages during embryogenesis does cell fate conversion occur. To address this question, we sought to perform a time course of endo labeling to determine the exact window of time at which CV ECs are lineage traced from endo progenitors.

First, we needed to obtain and characterize a Tamoxifen-inducible *Cre* line that was specific for endo cells. We found that *BmxCreERT2;Rosa26^tdTomato^* (Ehling et al. 2013) mice very specifically labeled endo and arterial cells when given Tamoxifen on E8.5 (Suppl. Fig. 1A). Arterial labeling should not influence the following experiments since previous studies have established that artery ECs are not CV progenitors (Mikawa and Fischman 1992). Immunostaining of E11.5 lineage-labeled hearts dosed at E8.5 and 9.5 (Fig. 1A) in tissue sections and whole mount preparations with the pan-endothelial markers Erg and Vegfr2 revealed recombination in greater than 90% of endo cells, but less than 3% of the SV (Fig. 1B-C; suppl. Fig. 1B). Across the timepoints, we noted that endo-derived CVs appeared first at E11.5 when they were exclusively in the septum (Fig. 1B’ and C). At E12.5 and E13.5, lineage-labeled CV ECs were mostly on the ventral myocardium where they appeared to progressively spread from the midline over time (Fig. 1D-top panel and suppl. Fig. 1C-top panel). Quantification showed that greater than 80% of ventral CV ECs were labeled while less than 5% of dorsal CV ECs were labeled (Fig.1E and suppl. Fig. 1C and D). Collectively, these patterns are consistent with previous studies and confirm the line’s utility in specifically labeling the endo lineage (Sharma et al. 2017a; Zhang et al. 2018).

**Figure 1.**
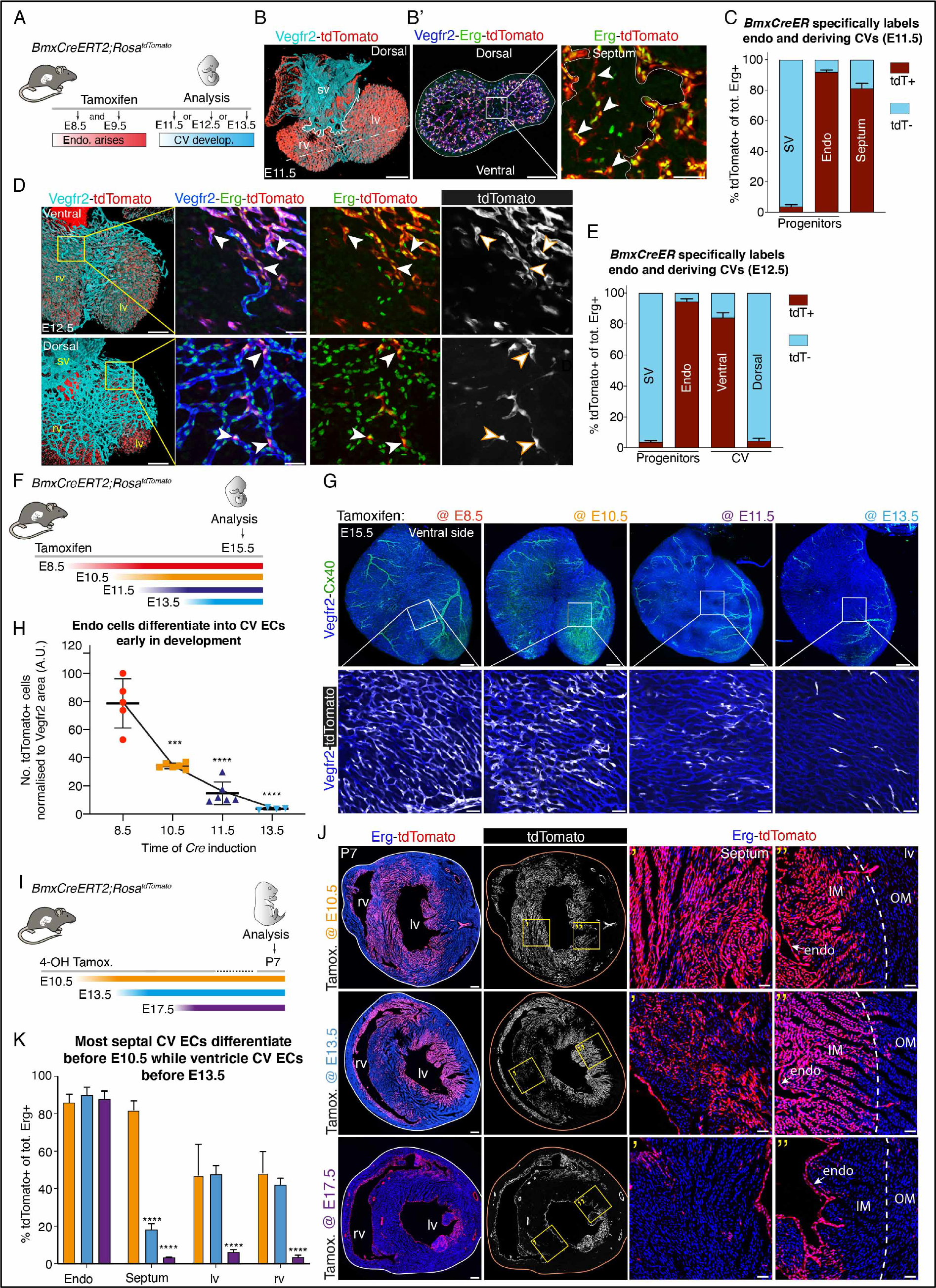
Endocardial-derived coronary vessels arose early in development, first in the septum then inner free walls. **A)** Experimental strategy for **B**-**E**. **B)** Confocal images showing whole-mount dorsal view of E11.5 *BmxCreERT2;Rosa^tdTomato^* lineage-labeled mouse heart (tdTomato in red; Vegfr2 in cyan). Solid line demarcates SV-derived ECs. Dotted line indicates location of section in **B’**. **B’)** Box highlights septal region and first endo-derived CVs (arrowheads). Vegfr2 (blue) and Erg (green) label endothelial cells (EC). **C)** Quantification of the total number of lineage-labeled tdTomato+;Erg+ cells from E11.5 *BmxCreERT2;Rosa^tdTomato^* heart sections (*n* = 4 hearts). **D)** Confocal images of E12.5 *BmxCreERT2;Rosa^tdTomato^* hearts. Boxed regions show the contribution of endo-derived CVs in the ventral and dorsal side (arrowheads). **E)** Quantification of the total number of lineage-labeled (tdTomato+;Erg+) endo and ECs in E12.5 *BmxCreERT2;Rosa^tdTomato^* whole-mount hearts (*n* = 6 hearts). **F)** Experimental strategy for **G** and **H**. **G)** Ventral whole-mount images of E15.5 *BmxCreERT2;Rosa^tdTomato^* hearts immunostained for Vegfr2 (blue) and the arterial marker Cx40 (green). Boxed regions show progressive reduction of endo-derived CVs (tdTomato+, white cells) from early vs. later *Cre* inductions. **H)** Quantification of the number of tdTomato+ cells per vessel area. Each symbols indicates a heart. **I)** Experimental strategy for **J** and **K**. **J)** Transverse heart sections from P7 lineage-traced *BmxCreERT2;Rosa^tdTomato^* mice immunostained for Erg (blue). Boxed regions show endo contributions to septal and left ventricle (lv) CVs (tdTomato+, red cells). Dotted lines indicate border between inner (IM) and outer myocardium (OM). **K)** Quantification of the number of lineage-labeled (tdTomato+;Erg+) cells in different regions of P7 hearts. E10.5, *n* = 6 hearts; E13.5, *n* = 4 hearts; E17.5, *n* = 4 hearts. Scale bar=200 μM in B, B’, D, G and J (full view). Scale bar=50 μM in B’, D, G, (boxed regions). Data are mean ± s.d. \*\*\**P* ≤ 0.001, ****, P ≤ 0.0001, by Student’s *t*-test.

We next labeled the endo at various stages and quantified lineage traced cells during development to establish when endo cells differentiated into CV ECs (Fig. 1F). For embryonic analysis, labeling was observed in CV ECs on the ventral side of the heart in whole mount preparations at E15.5. Surprisingly, there was very little contribution when endo cells were labeled after E11.5 (Fig 1G and H). The most robust contribution occurred with E8.5 labeling (Fig 1G and H). This was not due to reduced endo labeling with the later injection. Endo recombination was comparable in E15.5 heart sections induced at E9.5 and E12.5 (suppl. Fig.2A and B), while many more CV ECs were labeled with the early dose (suppl. Fig.2A and C). Thus, endo cells differentiated into CV ECs primarily during early heart development—mostly prior to E12.5.

**Figure 2.**
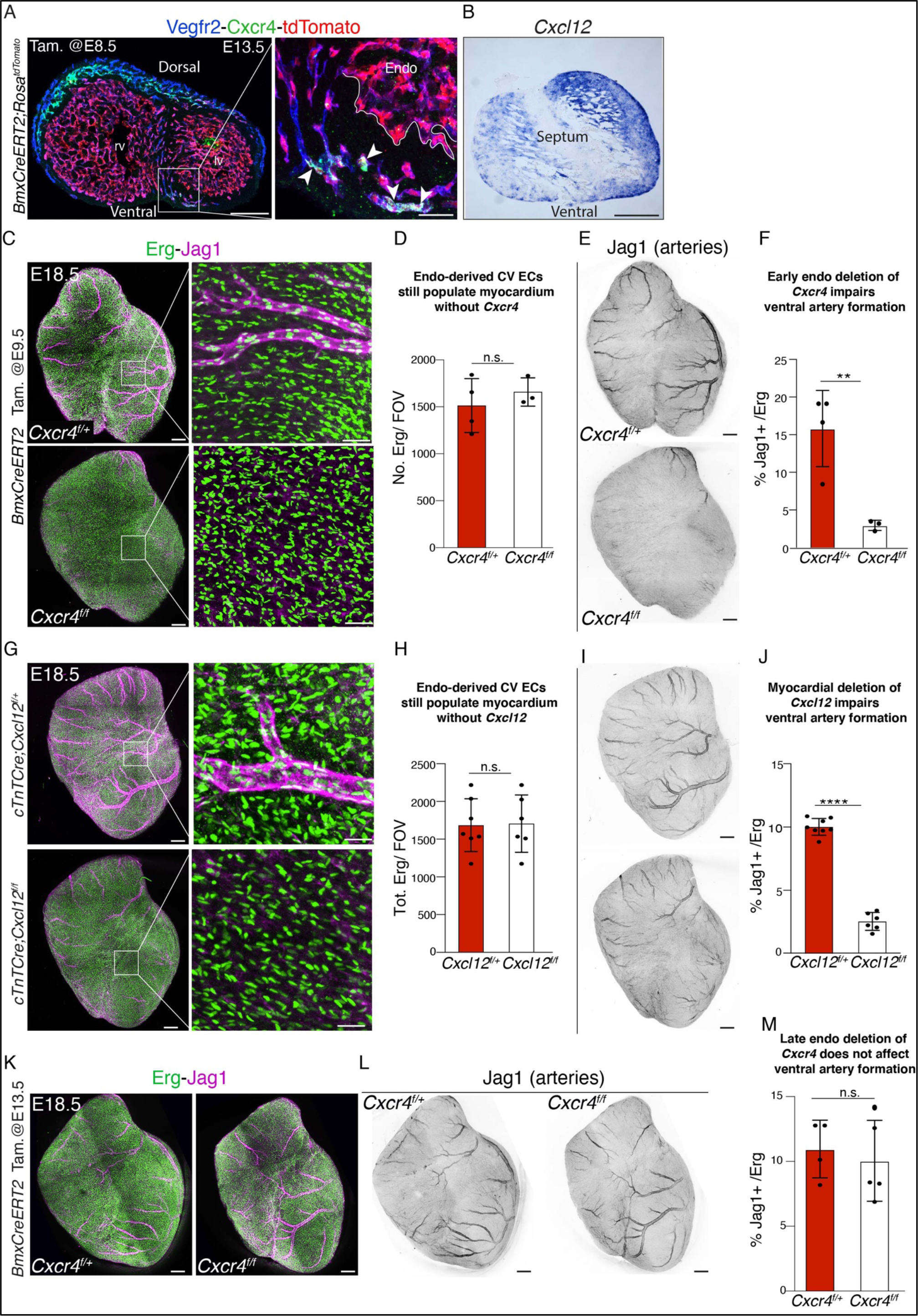
Cxcl12-Cxcr4 signaling was critical for artery formation rather than general endothelial migration. **A)** Immunostaining for Cxcr4 (green) and Vegfr2 (blue) in E14.5 *BmxCreERT2;Rosa^tdTomato^* lineage-traced heart section. Boxed region shows endo-derived CV expressing Cxcr4 (arrowheads) but lack of expression in endocardium (endo). **B)** *In* situ hybridization of *Cxcl12* in E14.5 WT heart section reveals strong expression in lv myocardium, which is attenuated in septum. **C** and **D)** Ventral view of E18.5 *BmxCreERT2*;*Cxcr4^f/+^* (controls, *n* = 4 hearts) and *BmxCreERT2;Cxcr4^f/f^* (mutants, *n* = 3 hearts) immunostained with Erg (green, all EC nuclei) and Jag1 (magenta, artery) upon early (E9.5) endocardial deletion of *Cxcr4*. Similar number of total ECs (Erg+ cells) is observed in control vs. mutant hearts indicating normal general migration, quantified in **D. E** and **F**) Ventral artery formation (Jag1+, black) is largely abrogated in *BmxCreERT2;Cxcr4^f/f^* embryos, quantified in **F**. **G** and **H)** Myocardial deletion of *Cxc12* recapitulates phenotype observed upon early endocardial deletion of *Cxcr4*. Quantification of total ECs number (Erg+ cells) from *cTnTCre;Cxcl12^f/+^* (controls, *n* = 6 hearts) and *cTnTCre;Cxcl12^f/f^* (mutants, *n* = 6 hearts) hearts indicate lack of gross migration defects. **(I** and **J)** Significant reduction of ventral artery formation (Jag1+ cells, black). **K-M)** Later (E13.5) endocardial deletion of *Cxcr4* does not affect arterial development. Scale bar=200 μM in A, B, C, E, G, I, K and L (full view). Scale bar=50 μM in A, C, and G (boxed regions). Data are mean ± s.d. \*\**P* ≤ 0.01, ****, P ≤ 0.0001, n.s., not significant, by Student’s *t*-test.

To ascertain whether the “second wave” of coronary expansion into the inner wall of the myocardium also arises from CV ECs that differentiated from endo early in development, we followed labeling at either E10.5, E13.5, or E17.5 out to postnatal day (P)7 (Fig 1I). With all dosing strategies, more than 80% of endo cells were labelled (Fig 1J and K). The E10.5 dose labeled approximately 80% of septal and more than 40% of the ventricular CV ECs while the E13.5 dose labeled much fewer septal cells but a similar number of those in the ventricle (Fig 1J and K). In contrast, the E17.5 dose resulted in very little labeling of CV ECs (Fig 1J and K). Note that the isolated labeling of coronary arteries at E17.5 is likely due to the expression of *Bmx* in arterial endothelial cells (Fig. 1J).

Our findings indicated that endo differentiation is spatiotemporally patterned and occurred almost exclusively during development. Most septal CV ECs arise during ventricular septation ∼between E8.5 and 10.5 and labeling dynamics suggest they are the precursors of ventral CV ECs. During later stages, shortly after E13.5, there is minimal cell fate conversion at the septum, but differentiation is still populating the inner myocardial wall/papillary muscle of the ventricles. Finally, levels of differentiation are very low, but still present (∼2.5%), in the inner myocardial wall/papillary muscle after E17.5. These data are in contrast with a previous model indicating that a second wave of CV ECs derives from postnatal endo differentiation (Tian et al. 2014), but consistent with a recent report demonstrating that this cell fate transition occurs mostly prior to E16.5 (Lu et al. 2021).

### Cxcl12/Cxcr4 deletion has a greater effect on artery formation than endo angiogenesis

The above data suggest that a prominent pathway from endo to CV ECs is a cell fate conversion at the septum followed by migration of septal CV ECs ventrally to populate the front of the heart. The signal(s) driving this specific migration event have not been described. Cxcl12 and Cxcr4 are good candidates since they stimulate CV ECs migration during Zebrafish coronary development (Harrison et al. 2015). However, Cavallaro *et al*. detected the opposite in mouse hearts, i.e. increased SV sprouting upon *Cxcl12* or *Cxcr4* knockout (Cavallero et al. 2015). We specifically investigated their role in early endo angiogenesis. We found that *Cxcr4* was expressed in endo-derived CV ECs at the junction between the septum and ventral myocardium, but not in the endo (Fig. 2A and suppl. Fig. 3A). Left ventricle cardiomyocytes expressed *Cxcl12* (Fig. 2B and Suppl. 3B). Thus, *Cxcr4* and *Cxcl12* are at the right time and place to drive migration of endo-derived CV ECs into ventral myocardium.

**Figure 3.**
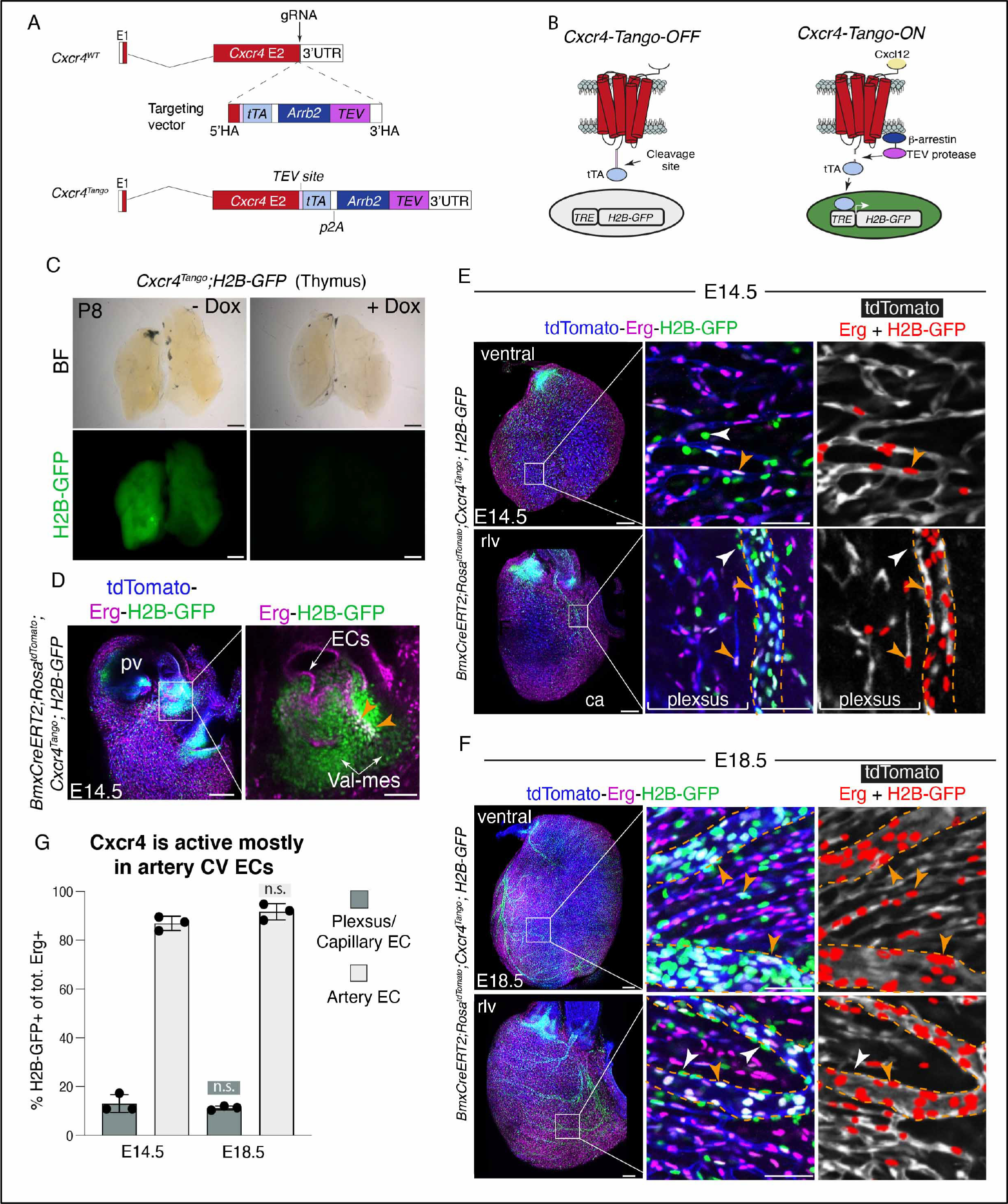
A novel Cxcr4 activity reporter revealed that most receptor activation is in arteries. **A** and **B)** Knock-in strategy used to generate *Cxcr4*-Tango allele, and schematic representation of activity. **C)** Brightfield (BF) and fluorescent images of P8 thymus from *Cxcr4^tango^;H2B-GFP* animals shows uniform H2B-GFP expression, indicating strong Cxcr4 activity. In doxycycline treated animals, H2B-GFP expression is not detected. **D)** E14.5 aortic and pulmonary valves (pv) view of *BmxCreERT2;Rosa^tdTomato^;Cxcr4^tango^;H2B-GFP* heart immunostained for Erg. Boxed region highlights Cxcr4 activity (H2B-GFP, green) in valve mesenchymal cells (val-mes). Orange arrowheads indicate Cxcr4 activity in ECs. **E** and **F)** Ventral and right lateral (rlv) view of E14.5 (**E**) and E18.5 (**F**) *BmxCreERT2;Rosa^tdTomato^;Cxcr4^tango^;H2B-GFP* hearts. Endo-derived CVs (tdTomato+) are shown in blue. High magnification views of boxed regions and colocalization analysis show few Cxcr4 active ECs within E14.5 vascular plexus and E18.5 capillary bed (orange arrowheads). ECs of coronary arteries (ca, dotted lines) show strong Cxcr4 activity. White arrows indicate Cxcr4 activity in pericytes/ smooth muscle cells (Erg-;H2B-GFP+). **G)** Quantification of endothelial Cxcr4 activity show higher percentage in arteries with no significant differences between the two ages (n=3 hearts stage). Scale bar=200 μM in C, D, E and F (full view). Scale bar=50 μM in D, E, and F (boxed regions). Data are mean ± s.d. n.s., not significant, by Student’s *t*-test.

Mouse genetics were used to test this hypothesis. Deletion of *Cxcr4* from endocardium using *BmxCreERT2* and early Tamoxifen dosing (E9.5) specifically knocked out the receptor in the endo- derived CV ECs (. Fig.3C). Surprisingly, Erg+ CV ECs still populated the ventral myocardium (Fig 2C and D). In contrast, artery differentiation was severely stunted. There was a four-fold decrease in Jag1+ arterial ECs on the ventral side of the heart (Fig. 2E and F) while those on the dorsal side, which are not subjected to *BmxCreER* deletion, remained intact (data not shown). Similar phenotypes were observed when *Cxcl12* was removed from cardiomyocytes (Fig. 2G-J). Timed Tamoxifen dosing confirmed our above results that the majority of endo to CV ECs differentiation occurs early in heart development. Specifically, deleting *Cxcr4* at E13.5 had no effect on either migration or artery formation (Fig. 2K-M). These data indicate that even though *Cxcr4* and *Cxcl12* are expressed where the vessels spread from the septum, migration is not dramatically affected by knocking down these genes. Instead, the primary defect is artery formation which is only seen when deletion in induced at E9.5.

These data suggested that observing the location of receptor activation might be a better predictor of Cxcr4 function than ligand-receptor expression patterns. But there was currently no available mouse line with this capability. We generated a new knock-in mouse line for this purpose by adapting elements of the TANGO system (Kono et al. 2014). CRISPR/Cas9 was used to insert, immediately after the *Cxcr4* stop codon, a bicistronic construct sequentially containing: 1) A tobacco etch virus (TEV) protease recognition site followed by a tetracycline transcriptional activator (tTA); 2) a P2A self-cleaving peptide sequence and 3) a mouse Beta-arrestin-2 fused to TEV protease (Fig. 3A). When this line is crossed with a tTA-activated nuclear GFP reporter (*H2B-GFP*)(Tumbar et al. 2004), ligand binding was designed to recruit B-arrestin-TEV, which cleaves the tTA, allowing it to move into the nucleus and stimulate H2B-GFP expression (Fig. 3B). Indeed, we observed H2B-GFP expression that was depleted with doxycycline (Dox) administration in multiple organs (Fig. 3C and suppl. Fig. 4). Thus, our genetic system performed as expected.

**Figure 4.**
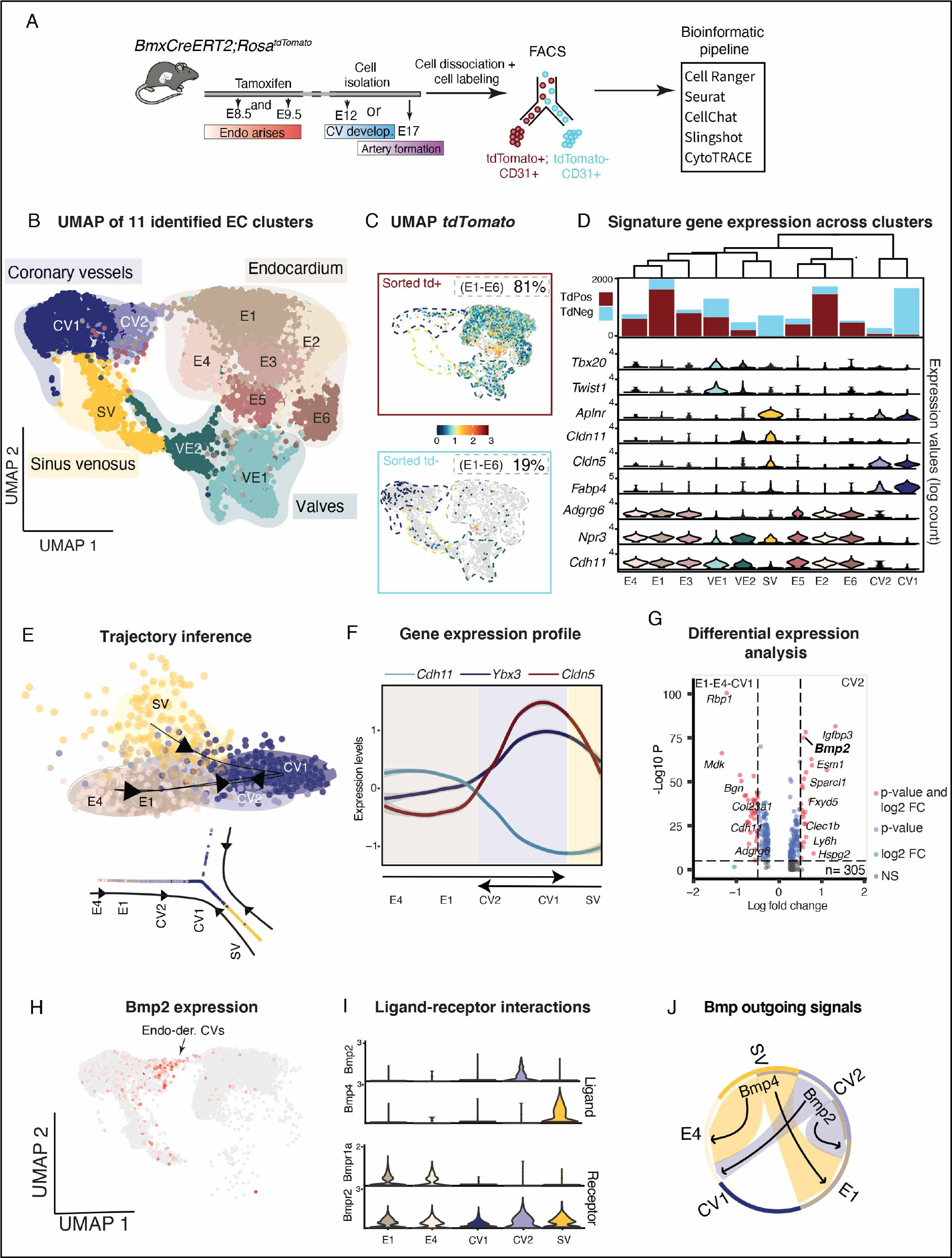
ScRNAseq of E12 cardiac ECs revealed upregulation of Bmp2 signaling during endo- to-CV transition. **A)** Overview of lineage tracing and scRNAseq experimental scheme and bioinformatics workflow. **B)** The Uniform Manifold Approximation and Projection (UMAP) plot of the 11 identified EC clusters grouped into 4 subtypes: Endo (E1-E6); Coronary vessels (CV1 and CV2); Sinus venosus (SV); Valve (VE1 and VE2). **C.** UMAP visualization of *tdTomato* gene expression in cells sorted as either *tdTomato+* (top panel) and *tdTomato-* (bottom panel). Colors of dotted lines indicate clusters identities as labeled in **B**. Percentages of total endo cells belonging to each category are indicated. **D)** Bar plots indicating ratios of *tdTomato+* and *tdTomato-* cells in each cluster. Clusters are ordered based on their relationship (similarity, top tree). Violin plots show expression of genes enriched in EC clusters used for cell subtype identification. **E)** Visualization of the trajectory inferred using slingshot on a dimensionality reduction of the cells (top panel) and as a 2D graph (bottom panel). **F)** Gene expression profile of 3 selected markers along the 2D trajectory from **E** (*y axis=cells*). **G)** Volcano plot showing differentially expressed genes between CV2 vs E1- E4 + CV1 clusters (cut-off for log2FC is >|2|; cut-off for P value is 10e-6). **H)** *Bmp2* expression is enriched in CVs transitioning from endo. **I)** Gene expression distribution of signaling genes related to Ligand-Receptors pairs of BMP signaling pathway. **J)** Chord plot showing all the significant interactions Ligand-Receptors pairs associated with BMP signaling pathway.

To characterize Cxcr4 activation in coronary vessels, *Cxcr4^tango^;H2B-GFP* hearts were immunostained with Vegfr2 and Erg. We observed many H2B-GFP-positive cells in lineage-labeled hearts, including valve mesenchyme (Fig. 3D). Only 10% of capillary plexus CV ECs expressed the Cxcr4 activation marker, including those on the ventral side lineage-labeled from the endo (Fig. 3E- G). Consistent with a specific role for Cxcr4 in artery development (see Fig. 2E and F), approximately 90% of artery ECs expressed H2B-GFP in both stages analyzed (Fig. 3E-G). Interestingly, perivascular cells, likely pericytes and smooth muscle cells were also positive (Fig. 3E and F). These data establish the validity of a new tool for probing Cxcr4 activation and highlights how activation reporters can be better predictors of loss-of-function phenotypes than ligand-receptor expression.

### ScRNAseq reveals a *Bmp2*+ transitioning population during early coronary development

Since Cxcl12/Cxcr4 signaling was critical at a later than expected stage during endo angiogenesis, i.e. artery differentiation, we next sought to discover the pathways involved in earlier steps in the process. scRNASeq was performed on endo and CV ECs from E12 *BmxCreERT2;Rosa26^tdTomato^* -lineage labeled hearts, which is the stage shortly after the onset of endo differentiation. Cells were FACS-sorted into CD31+/tdTomato+ (endo and its derivatives) and CD31+/tdTomato- (SV and its derivatives) populations and processed using the 10X Genomics platform (Fig. 4A). After quality control and filtering (Suppl. Fig. 5A-D and Methods), a total of 10,878 cells were available for cell type characterization. Unbiased clustering and uniform manifold approximation and projection (UMAP) analysis revealed 11 cell clusters (Fig. 4B). Cell type identities were assigned using known gene expression. Six clusters expressed the endo markers *Cdh11*, *Npr3,* and *Adgrg6* and were mostly *tdTomato+* (Fig. 4C and D). Two clusters of CV ECs expressed the capillary markers *Cld5* and *Fabp4* (Fig. 4C and D). Only a small fraction of *tdTomato+* cells were found in the CV EC clusters at this stage, indicating that most were derived from the SV (Fig. 4C and D). We also detected two clusters of valve endothelial cells (*Twist1+*;*Tbx20+*) containing approximately the same number of *tdTomato* positive and negative cells (Fig. 4C and D). These cells and lineage identities are consistent with our previous data and the above lineage tracing, validating the dataset for further analysis.

We next subsetted the data to increase the resolution of our CV angiogenesis analyses. The endo clusters closest to CV ECs in the UMAP (E1 and E4), the SV cluster, and the CV clusters (CV1 and CV2) were subjected to trajectory analysis using Slingshot (Street et al. 2018), which infers cell state transitions. SV cells were predicted to transition into CV1 while endo cells transitioned first into CV2—the CV cluster also positive for the lineage marker—and then CV1 (Fig. 4E and Suppl. Fig. 6A). We next plotted the 50 most relevant genes reconstructing these cell state transitions (Suppl. Fig. 6B). Along the trajectory E4-E1-CV2, we observed a down-regulation of endo-defining genes (*Cdh11; Npr3; Irx5*) as the cells became CV ECs (*Cldn5; Fabp4*). This was accompanied by an induction of protein synthesis and metabolic related genes (*Ybx3; Eif5a; Aldoa*) (Fig. 4F and Suppl. Fig. 6B).

Differential gene expression (DEG) values comparing CV2 to aggregated E1, E4, and CV1 were calculated to search for genes that might have a specific role in endo angiogenesis. We found up-regulation of Vegf-A-induced genes (*Igfbp3*; *Esm1*) in CV2 (Fig. 4G), which is consistent with previous data demonstrating that endo sprouting involves Vegf-A while SV sprouting is instead stimulated by Vegf-C and Ela (Wu et al. 2012; Chen et al. 2014a; Sharma et al. 2017b). During DEG analysis, *Bmp2* stood out as being particularly specific to CV2 (Fig. 4G and 4H). Bmp2 is an attractive candidate because it’s known to dramatically affect endo fate during valvulogenesis (Ma et al. 2005). Overall, previous studies showed that Bmp2 induces endo ECs to transition into valve mesenchyme by stimulating the endothelial-mesenchymal transition (endMT)(Rivera-Feliciano and Tabin 2006; Luna-Zurita et al. 2010), and *in vivo Bmp2* overexpression inhibits endo genes, driving cells into a mesenchymal state (Papoutsi et al. 2018; Prados et al. 2018). We therefore performed additional analysis of Bmp ligand and receptor signaling pathway genes.

A ligand-receptor interaction analysis tool, CellChat, identified the *Bmp2* ligand as being specifically expressed in CV2 while its receptor subunit *Bmpr1a* was specifically in endo cells (Fig. 4I). The complementary receptor subunit, *Bmpr2,* (Bmps signal through heterotetramers of two receptor subunits) was in all clusters analyzed (Fig. 4I). The other Bmp ligand detected was *Bmp4*, which was highly specific to SV sprouts (Fig 4I). A Chord plot representing Bmp interactions predicted Bmp2 from CV2 and Bmp4 from SV signaling to a subset of endo ECs within E1 and E4 (Fig. 4J). In addition to Bmp signaling, CellChat also detected additional outgoing signals from endo cells to the CV2 cluster, including Vegf and Notch signaling pathways, but a Cxcl12/Cxcr4 relationship was not detected during this early stage (Suppl. Fig. 6H).

In total, our trajectory analysis using scRNAseq with lineage labeling information indicated that a cluster of newly formed CVs expressing *Bmp2* signals to the adjacent endo, possibly to promote endo-to-CV ECs differentiation and/or angiogenesis. This model was reinforced by the results obtained using CytoTRACE (Gulati et al. 2020), a trajectory analysis tool that uses the number of genes expressed to assign differentiation state. Here, an endo transition through CV1 and then CV2 was seen where the CV2 “bridge” to CV1 was enriched for Bmp2 (Suppl Fig. 6C-G).

**Figure 5.**
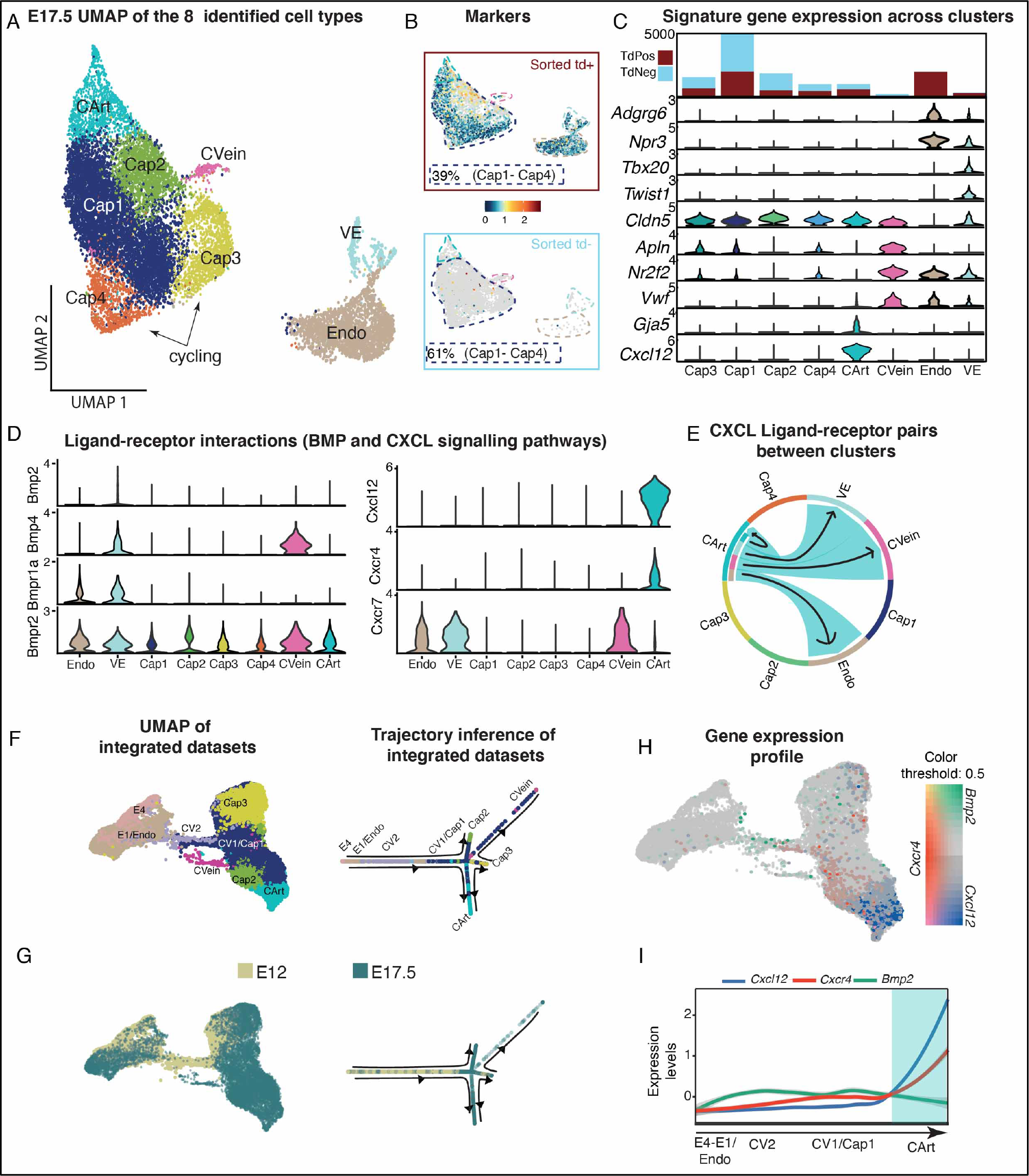
Combined E12 and E17.5 scRNAseq indicated the endo-to-CV transition and Bmp2 expression is largely restricted to the early time point. **A)** The Uniform Manifold Approximation and Projection (UMAP) plot of the 8 identified EC clusters from E17.5 grouped into 5 subtypes: Endo; Valve (VE); Coronary Capillary (Cap1-Cap4); Coronary Veins (CVein); Coronary Arteries (CArt). **B)** UMAP visualization of *tdTomato* gene expression in cells sorted as either *tdTomato+* (top panel) or *tdTomato-* (bottom panel). Colors of dotted lines indicate clusters identities as labeled in **A**. Percentages of total capillary cells belonging to each category are indicated. **C**) Bar plot indicating ratios of *tdTomato+* and *tdTomato-* cells in each cluster. Clusters are ordered based on their relationship (similarity, top tree). Violin plots show expression of genes enriched in EC clusters used for cell subtype identification. **D** and **E)** UMAP of E12/17.5 integrated dataset (left panel) and visualization of the trajectory as a 2D graph structure (right panel), colored according to clusters of origin (**D**) or developmental stage (**E**). **F**) UMAP of integrated dataset colored by expression of the 3 indicated genes. **G)** Gene expression profile of 3 markers along the 2D trajectory of integrated dataset (y axis cells). **H)** Gene expression distribution of signaling genes in E17.5 dataset related to Ligand-Receptors pairs of BMP (left) and CXCL (right) signaling pathways. Bmp2 is down at this stage and Cxcl12 is up. I) Chord plot showing all the significant interactions (L-R pairs) associated with CXCL signaling pathway.

**Figure 6.**
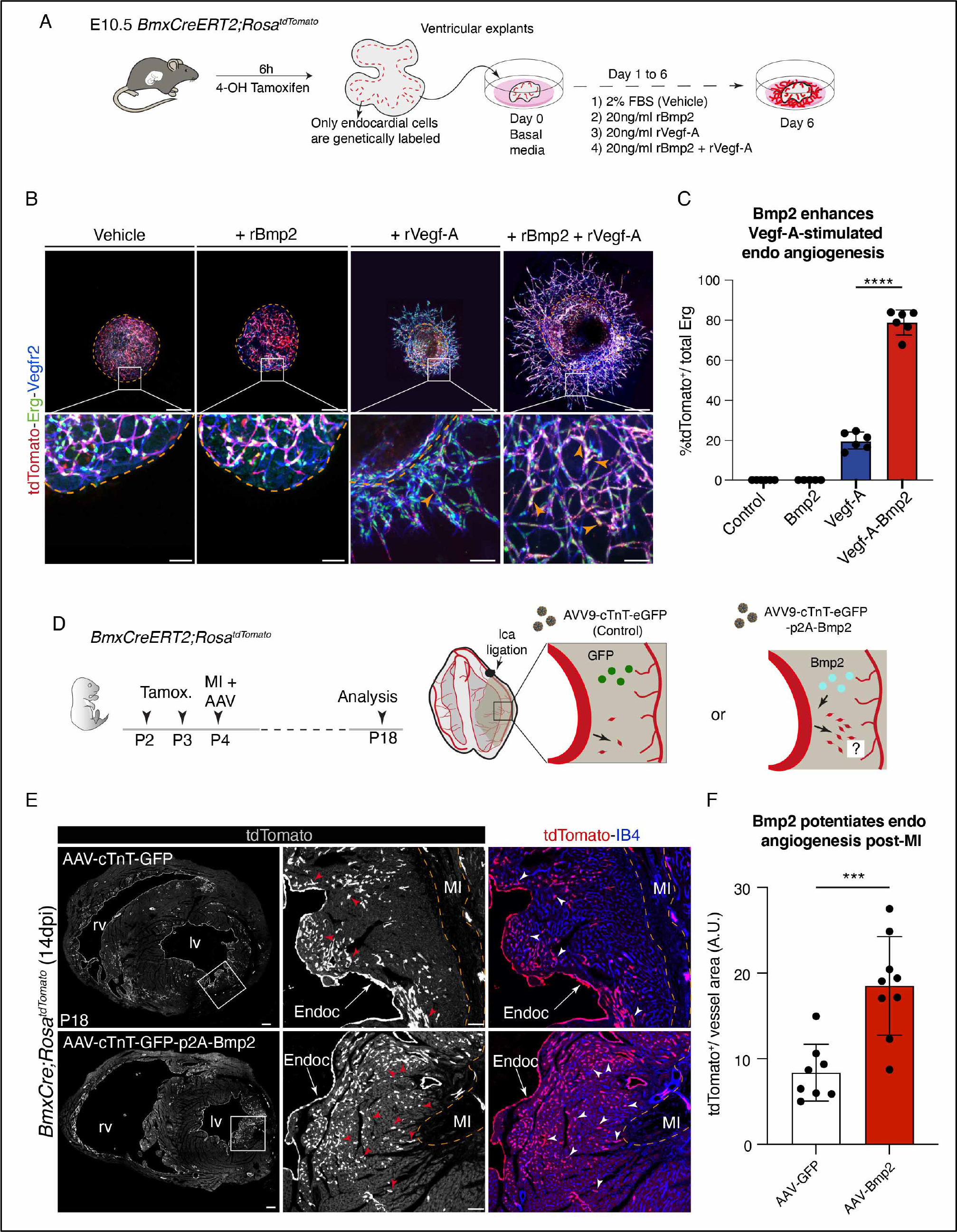
Bmp2 potentiates endo-derived CV angiogenesis in culture and neonatal hearts. **A)** Experimental strategy of E10.5 ventricular explants treated with recombinant Bmp2. **B)** Immunostaining for Erg (green) and Vegfr2 (blue) in E10.5 lineage labeled ventricular explants. Boxed regions show no ECs sprouting in explants treated with 2% FBS (vehicle) or 20ng/ml rBmp2. Combined Vegfa and Bmp2 treatment (rightmost panel) strongly induces endo-derived CVs sprouting (tdTomato+ cells, arrowheads). **C)** Quantification of total number of endo-derived CVs (tdTomato+ in sprouting ECs) from E10.5 ventricular explants shows increased endo-derived CVs sprouting in explants treated with combination of Vegfa and Bmp2. (rBmp2, *n* = 5 explants; Vegfa, *n* = 6 explants; combination, *n* = 6 explants) **D)** Experimental strategy of neonatal left coronary artery (lca) ligation on lineage traced hearts injected with cardiac specific AAVs expressing GFP (AAV- cTnT-GFP, vehicle) or Bmp2 (AAV-cTnT-GFP-p2A-Bmp2) in **E** and **F**. **E)** P18 tissue sections from *BmxCreERT2;Rosa^tdTomato^* injured hearts stained for tdTomato (red) and isolectin B4 (IB4, blue). Boxed regions show increased endo-derived CVs (arrowheads) in the left ventricles injected with AAV expressing Bmp2 injected hearts (*n* = 9 hearts, lower panels) compared with vehicle-treated hearts (n=8, upper panels). **F)** Quantification of total number of tdTomato*+* CVs per vessel area. Scale bar=200 μM in B, and E (full view). Scale bar=50 μM in B, and E (boxed regions). Data are mean ± s.d. ***P ≤ 0.001, ****, P ≤ 0.0001, by Student’s t-test.

### Late gestation scRNAseq lacks a prominent transitioning population

Because staged lineage analysis revealed minimal endo-to-CV EC differentiation after E17.5 (see Fig. 1J and K), we hypothesized that critical cell states and genes could be revealed by comparing early and late scRNAseq datasets. E17.5 hearts were harvested and subjected to the same pipeline as above (Fig. 4A), resulting in high quality data from 13,454 cells (Suppl. 5E-H). Gene expression identified CV ECs, endo cells, and valve cells, which separated into a total of 8 clusters (Fig. 5A). The endo cluster was primarily *tdTomato+* while approximately half of capillaries and arteries expressed the lineage label (Fig. 5B) consistent with previous lineage analysis. Veins were mostly *tdTomato-*, which is expected giving their localization to mostly SV-derived regions (Fig. 5B and C)(Chen et al. 2014a). The most diverse population in the dataset was CV ECs, which clustered into the following subpopulations: 4 *Cldn5+, Apln+* capillary clusters (Cap1 and Cap2= non cycling ECs and Cap2 -Cap3= cycling ECs), a *Gja5+,Cxcl12+* arterial cluster, and a *Vwf+* vein cluster (Fig. 5C). In contrast to the E12 dataset, there was no population bridging endo cells and CV ECs in the UMAP (compare Fig. 5A with Fig. 4B), a result also observed using CytoTRACE (Suppl. Fig. 7A). These data agreed with lineage tracing experiments (Fig. 1F-K and (Lu et al. 2021)), further indicating that most endo-to-CV EC fate transitions occur before E17.5.

Next, data from both time points were integrated to further explore the potential absence of a transitioning population at E17.5. After integration, 10 clusters were detected and annotated based on their nomenclature in the single datasets (Fig.5D and Suppl Fig. 8A). Endo cells across the two datasets mostly overlapped, suggesting that the endo cell state may not dramatically change over developmental time (Fig. 5D and E; Suppl Fig. 8B). Similarly, capillary clusters CV1 and Cap1 from E12 and E17.5, respectively, were also largely indistinguishable. In contrast, cells within CV2 from the E12 sample were clustered apart from the rest of E17.5 capillary clusters (Fig. 5D and E; Suppl Fig. 8C). The UMAP plot and trajectory analysis showed that CV2 cells were unique to E12, and that they existed in a transcriptional state between endo and CV1/Cap (Fig. 5D and E). These data further suggest that CV2 is an immature transitioning CV ECs population only present at the earlier developmental stage.

Integrated data also confirmed that *Bmp2* was specific to E12 CV2 (the putative transition population), and the *Cxcl12/Cxcr4* axis specific to later artery populations (Fig. 5F and G). CellChat did not predict Bmp2 signaling from CV ECs to endo ECs at E17.5 as it did for E12 (Fig. 5H and Suppl. Fig. 7D). However, Bmp4 was still predicted to be an autocrine signal within the vein compartment (Fig. 5H), which is consistent with previous studies demonstrating BMP as the vein specification factor early in development (Neal et al. 2019). Interrogation of the Cxcl pathway predicted autocrine Cxcl12/Cxcr4 signaling within the arterial cluster. Interestingly, the Cxcl12 sink receptor *Cxcr7* was expressed in Endo, VE, and Vein clusters, suggesting a possible role for opposing arterial differentiation in veins (Fig. 5H and I). In total, *in-silico*-predicted ligand-receptor pairs support our findings that Cxcl12/Cxcr4 signaling in the developing heart is most critical for artery development, and that Bmp2 might be a signal stimulating the endo-to-CV EC transition, specifically at early stages.

### Exogenous Bmp2 facilitates Vegf-A-induced endo angiogenesis

To test our hypothesis that Bmp2 induces early endo angiogenesis, we performed an *in vitro* coronary sprouting assay using lineage labeled E10.5 heart ventricle explants cultured on Matrigel for 6 days with different combinations of recombinant proteins (Fig. 6A). There was no sprouting of Vegfr2-positive vessels in wells containing either control vehicles or Bmp2 alone (Fig. 6B and C). At the 20ng/ml concentration, Vegf-A stimulated sprouting, but only 20% of vessels were tdTomato- positive, indicating that most did not derive from the endo (Fig. 6B and C). In contrast, explants treated with both Vegf-A and Bmp2 exhibited robust sprouting, most of which involved endo-derived ECs (Fig. 6B and C). These data demonstrated that, at least in this assay, Bmp2 alone does not induce CV EC outgrowth, but it does greatly enhance Vegf-A-stimulated endo angiogenesis.

We next investigated if Bmp2 could enhance endo angiogenesis in the context of injury. *BmxCreERT2;Rosa^tdTomato^* neonates were injected with Tamoxifen and 2 days later subjected to MI through a permanent left coronary artery (LCA) ligation. To test the effect of locally delivered Bmp2, we generated an adeno-associated viral vectors expressing either Bmp2 (AAV-Bmp2) or GFP (control, AAV-GFP) under the control of the cardiac troponin T (cTnT) promoter. AVVs were administrated immediately after MI (Fig. 6D). One week post-MI, transduced hearts already showed widespread GFP expression in cardiomyocytes and qualitatively increased angiogenesis from the endo in Bmp2-treated samples (Suppl. Fig. 9). Quantification at two weeks post-MI, showed that AAV-Bmp2 hearts had more than a twofold increase in endo-derived CV ECs adjacent to the endo (Fig. 6E-F). This was in contrast with non-injured hearts where Bmp2 did not have the same effect (data not shown). Since Vegf-A is expected to be upregulated upon MI, the difference in AAV- Bmp2’s effect in non-injured and MI is similar to explants where Vegf-A is required for Bmp2’s activity. These data, coupled with the observation that Bmp2 is specific to a transitioning population, supports the model that Bmp2 is involved in stimulating early endocardial angiogenesis, potentially through new sprouts signaling to endo cells.

## DISCUSSION

Endocardial cells that line the heart chambers give rise to multiple tissues in the developing heart, including CV ECs, CV mural cells, and valve mesenchyme (Zhang et al. 2018). Its intrinsic plasticity may offer a valid cell-based therapeutic approach to ameliorate pathological heart conditions. Although the contribution of the endo-to-CVs has been extensively studied and debated, the molecular mechanism triggering this transition and whether endo retains its angiogenic potential throughout development remains largely unknown.

Our studies utilizing time resolved lineage tracing provide a greater understanding of when endo-derived CVs originate and how they populate the heart (Fig. 7A). Most endo-derived CVs originate at the onset of septal development when endo cells differentiate into septal CV ECs. Subsequently, these endo-derived CVs migrate ventrally from the septum into the intramyocardial space. This pathway is supported by two observations: 1. Few septal and ventral CVs are lineage traced when endo is labeled after E11.5 and 2. At this time point, there are only endo-derived CVs in the septum. A second event occurs slightly later—until approximate E13.5—where endo cells differentiate and migrate laterally directly into the ventricular free walls. This conclusion is supported by lineage labeling in CVs of the free walls in the absence of sepal and ventral labeling with an E13.5 Tamoxifen injection. The fact that there is very little CV tracing with an E17.5 label indicates that most endo differentiation occurs during these earlier events, although the small amount of remaining tracing indicates some residual differentiation capabilities. These findings indicating early endo differentiation were supported by the absence of an apparent endo-to-CV transitioning population in our E17.5 scRNAseq data and the lack of phenotype when *Cxcr4* was deleted after E11.5. Together, the emerging model is consistent with and adds significant spatiotemporal details to recent findings that little, if any, endocardial differentiation occurs after birth (Lu et al. 2021).

**Figure 7.**
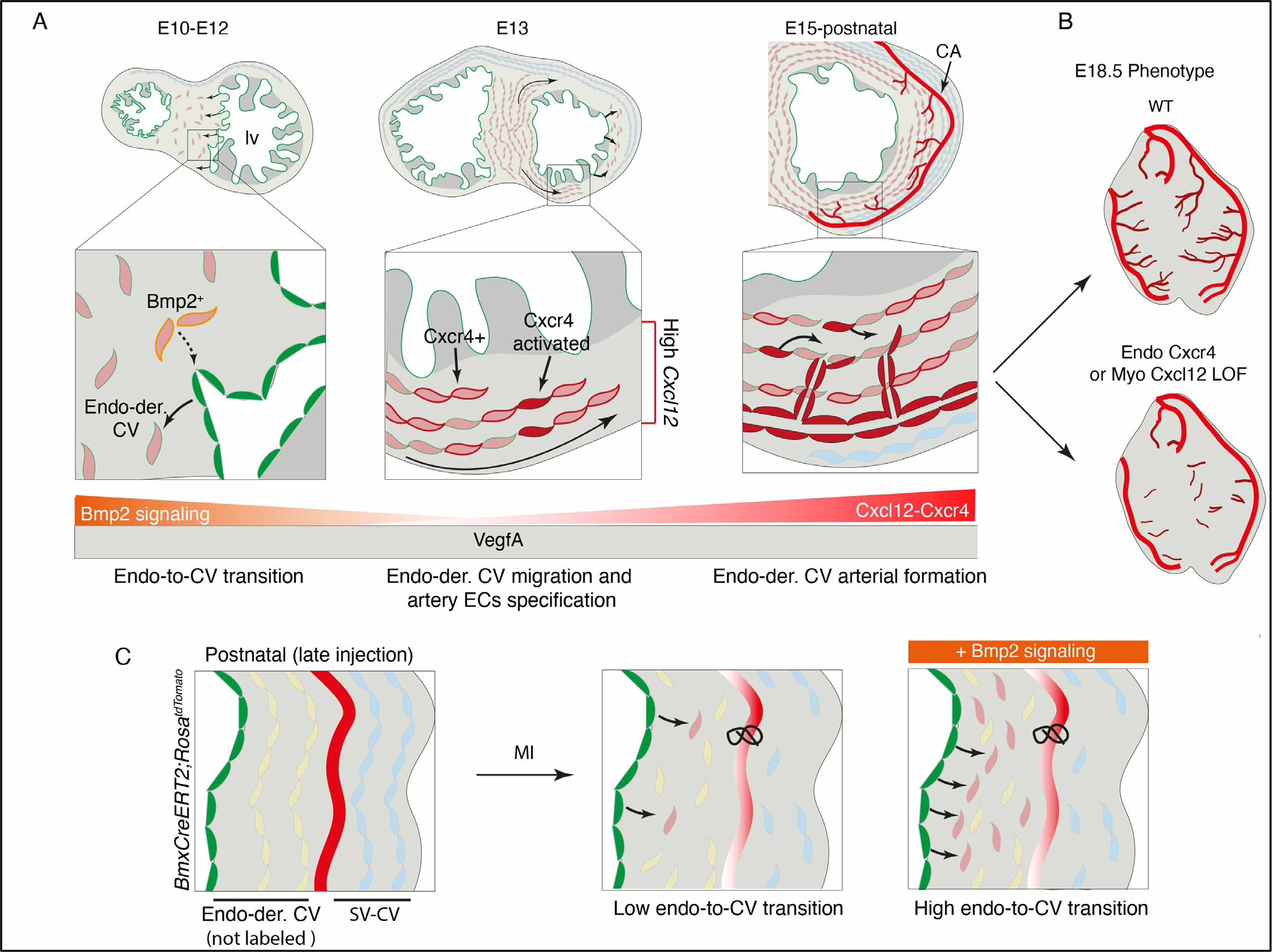
Sequential Bmp2-Cxcr4 signaling drives endo-to-CV ECs transition and artery formation during heart development and regeneration. **A)** Endo cells (green cells) differentiate into CV ECs (light red cells) within the developing septum. ECs expressing *Bmp2* signal to adjacent endo (dotted arrow) to favor endo-to-CV ECs transition. As development precedes, septal endo-derived CV ECs migrate ventrally into myocardial free wall. Cardiomyocytes expressing *Cxcl12* activate a subset of Cxcr4+ endo-derived CV ECs (red cells). Cxcr4 activated cells differentiate into artery promoting ventral arterial formation. **B)** Early endo *Cxcr4* deletion or systemic abrogation of myocardial *Cxcl12* disrupts endo-derived artery formation in the ventral side of the heart. **C)** Exogenous administration of Bmp2 greatly enhances endo-to-CV ECs transition in subendocardial space in neonatal mice subjected to experimental MI.

Cxcl12-Cxcr4 signaling has been previously implicated in coronary artery maturation in mice, which was a result of ligand expression around the outflow tract attracting CV ECs to the aorta during stem formation and blood flow initiation (Cavallero et al. 2015; Ivins et al. 2015). However, the role of Cxcl12 in endo-derived CV development at distances far from the outflow tract remained unexplored. Our lineage tracing and loss-of-function data demonstrate that Cxcl12-Cxcr4 signaling is critical for ventral arteries to develop from endo-derived CVs migrating from the septum to the ventral side of the heart (Fig. 7B).

Our data utilizing a novel mouse line identified a role specifically in artery formation also highlights the importance of considering the separation of gene expression and receptor activity. Expression of the chemokine *Cxcl12* and its receptor *Cxcr4* suggested that this signaling pathway may trigger the aforementioned septal-to-ventral migration of endo-derived CV ECs. However, genetic loss-of-function experiments revealed that deletion of *Cxcr4* from endo-derived CVs did not dramatically affect EC migration. Instead, *Cxcr4* deletion specifically abrogated artery formation on the ventral side of the heart where endo-derived CVs are highly concentrated.

These data underscored the importance of identifying the exact cells responding to a ligand within the tissue. To establish how endo-derived CVs expressing *Cxcr4* are activated during arterial remodeling, we generated a new mouse model to monitor Cxcr4 activity at the single cell level using the TANGO system (Kono et al. 2014). In this approach, Cxcr4 is engineered to contain a TEV protease site followed by a tTA transcriptional activator. When coupled with a *TRE-H2B-GFP* reporter, nuclear GFP is a readout of Cxcr4 activity. This tool showed that a small fraction of ECs in the vascular plexus are activated whereas strong Cxcr4 activity is found in forming arteries. This result supports the genetic phenotypes and indicate that only a subset *Cxcr4* expressing cells are activated by the ligand to function during artery remodeling, despite widespread *Cxcl12* expression in the region (Fig. 7A). The ability to record Cxcr4 activity *in vivo* will therefore represent a key tool to define how the secreted ligand finds and activates its receptors within the tissue during arterial development and regeneration.

Our demonstration that the endo’s natural angiogenic potential is largely restricted to early development is in agreement with a recent study indicating that the expansion of endo-derived CVs in the perinatal period is driven by angiogenesis of pre-existing vessels, especially at the inner myocardial wall (Lu et al. 2021). Therefore, these authors and our data provide evidence for an alternative interpretation of the previous model stating that *de novo* formation of postnatal CVs arise from endo differentiation occurring after birth (Tian et al. 2014). An outstanding question is how/why do embryonic endo cells lose their angiogenic capacity? Here, we analyzed early and late scRNAseq from lineage-traced ECs to begin to address this question. At the beginning of CV development (E12), we detected a subset of cells forming a transcriptional “transition” state between the endo and CV EC state, which was supported by overlying lineage information within the data onto trajectory analyses. Integrating the E12 and E17.5 datasets revealed that endo cells are transcriptionally similar across development, but that only E12 possess a “transitioning” population that specifically expresses *Bmp2*. These data suggest that stage specific Bmp2 could be involved in the restricted angiogenic period during development through a feed forward-like mechanism where new CV ECs signal to endo to stimulate more differentiation (Fig 7A).

*In vitro* and *in vivo* experiments supported this hypothesis. Recombinant Bmp2 potentiated VegfA-stimulated endo sprouting from embryonic heart explants. Adenoviral-mediated *Bmp2* overexpression in CM at the time of MI strongly enhanced the endo-to-CV transition in neonatal hearts (Fig. 7C). Other studies observed that ectopic *Bmp2* expression throughout heart development induced uncontrolled EMT within the ventricular endocardium resulting in an LVNC-like phenotype (Prados et al. 2018). In contrast, Bmp2 overexpression in neonatal hearts in our experiments did not produce histologically obvious ventricular phenotypes, likely because of the short time window and much later stage. Parallels with the embryonic study do exist. There, overexpressed *Bmp2* caused endo cells to downregulate endo marker genes, which occurs when endo cells are differentiating into CV ECs (Papoutsi et al. 2018; Prados et al. 2018).

Future studies should address whether Bmp2 can reactivate endo differentiation in adult hearts, and whether this improves recovery from cardiac injury. Endo angiogenesis has been demonstrated in adult injured hearts overexpressing Vegf-B in CM (Räsänen et al. 2021) or Vegfr2 in endo cells (Jiang et al. 2021), supporting the idea that adult endo cells do have the potential to differentiate. In total, our findings shed light on the molecular mechanisms orchestrating the cell fate conversion from endo-to-CVs and their role in arterial remodeling. We used this knowledge to promote the endo-to- CVs transition in injured neonatal hearts.

## METHODS

### Mouse Strain

All mouse husbandry and experiments were performed in accordance with Stanford Institutional Animal Care and Use Committee (IACUC) guidelines. Mouse lines used in this study were: wild type (CD1, Charles River Laboratories, Strain Code #022)*, Cxcl12^fl/fl^* (The Jackson Laboratory, B6(FVB)- *Cxcl12^tm1.1Link^*/J, Stock# 021773), *Cxcr4^fl/fl^* (The Jackson Laboratory, B6.129P2-*Cxcr4^tm2Yzo^*/J, Stock# 008767), *Rosa26^tdTomato^* Cre reporter (The Jackson Laboratory, B6.Cg-*Gt(ROSA)26Sor^tm9(CAG-TdTomato)Hze^*/J, Stock# 007909), *Rosa26^ntdTomato^* Cre reporter (The Jackson Laboratory, B6.Cg-Gt(ROSA)26Sortm75.1^(CAG-tdTomato*)Hze^/J, Stock# 025106)*, Cxcl12-DsRed* (The Jackson Laboratory, *Cxcl12^tm2.1Sjm^*/J, Stock# 022458), *pTRE-H2BGFP* (The Jackson *Tg^(tetO-HIST1H2BJ/GFP)47Efu^*/J, Stock# 005104), *BmxCreER* (Ehling et al. 2013). All mice in this study were maintained on a mixed background.

### Generation of a Cxcr4 Tango Knock-in allele

For the generation of Cxcr4-Tango we inserted Tango elements into the Cxcr4 endogenous locus by CRISPR/Cas9-mediated genome editing in C57BL/6J strain. Briefly: three guide RNAs overlapping the *Cxcr4* stop codon were selected for activity testing. Functional testing was performed by transfecting a mouse embryonic fibroblast cell line (MEF) with guide RNA and HiFiCas9 protein (Integrated DNA Technologies). Following transfection, the guide RNA target site was PCR amplified from transfected cells and analyzed by ICE (Synthego) to detect Cas9-mediated mutations. The guide RNA selected for genome editing in embryos was Cxcr4-g55B (protospacer sequence 5’- GTCTTTGCATAAGTGTTAGC-3’). 910 bp 5’ and 1014 bp 3’ homology arms were cloned into a plasmid donor vector flanking the Tango elements: 1) a codon-optimized tTA with preceding TEV protease cleavage site; 2) a P2A “self-cleaving” peptide sequence; 3) a Beta-arrestin-2-TEV protease fusion protein; 4) 3X stop codons, FRT site, and loxP-flanked SV40 late polyadenylation sequences. C57BL/6J zygotes were microinjected with 800 nM HiFiCas9 protein, 50 ng/ul guide RNA and 20 ng/ul supercoiled donor vector plasmid. Injected embryos were implanted in recipient pseudopregnant females. Eight resulting pups were screened by PCR for the presence of the knock-in allele. Two of 8 pups were positive for the knock-in event with primers spanning from outside of the homology arms to unique sequences in the knock-in allele. Both founders also showed evidence of vector backbone integration, consistent with the presence of a tandem integration event at the Cxcr4 target site. Founder animals were mated to C57BL/6J-Rosa26-CAG-Flpo transgenic animals to collapse tandem integration events to single-copy knock-ins. F1 animals heterozygous for the correct knock-in event and absence of vector backbone sequence were used for subsequent breeding to establish the knock-in colony. Details of genotyping will be provided on request.

### Breeding, tamoxifen administration and doxycycline treatment

Timed pregnancies were determined by the morning day on which vaginal plug was found as E0.5. To activate inducible *Cre*, 4mg of 20mg/ml Tamoxifen (Sigma-Aldrich, T5648) was administered to pregnant dams by oral gavage at E8.5 and E9.5. Two consecutive Tamoxifen doses were used to characterize the recombination rate of *BmxCreER* mouse line and for scRNAseq experiments. For time-course lineage tracing analysis and loss of function (LOF) experiments one dose of 4mg of Tamoxifen or 2mg of 4-OH tamoxifen (Sigma-Aldrich, H6278) was administrated at defined embryonic stage. For neonatal studies, two consecutive doses of 4mg Tamoxifen were administrated by oral gavage to nursing mom. In order to test the ability of doxycycline to effectively abolish H2B transcription mice were given food containing 200 mg of doxycycline per kg of diet (Bio-serv # S3888). Mice were given doxycycline containing food for six days prior to sacrifice and harvest of organs.

### Whole-mount immunofluorescence

All embryos were dissected in cold 1X PBS and fixed in 4% paraformaldehyde (PFA) at 4 °C with shaking for 1 h. Embryos were then washed twice (10 min each wash) with PBS at room temperature with shaking before dissection for whole-mount immunostaining. Whole hearts were washed in 0.5% PBT (1X PBS with 0.5% -Triton-X 100) at room temperature for one hour before incubation with primary antibodies. Primary antibodies were dissolved in blocking solution consisting of 5% donkey serum in 0.5% PBT. Hearts were incubated in the solution with primary antibodies with shaking overnight at 4 °C. Hearts were then washed every hour with 0.5% PBT for six hours with shaking at room temperature. Hearts were then stained with corresponding fluorescent-dye-conjugated secondary antibody secondary antibodies with the same conditions and procedure as for primary antibodies. After washing off the secondary antibodies with several washes of 0.5% PBT, hearts were transferred in 50% glycerol for 1 h and then mounted in anti-fade mounting media solution containing 100% glycerol + 20%(w/v) n-propyl gallate (Sigma P3130).

Imaging was performed using Zeiss LSM-700 confocal microscope (10× or 20× objective lens) with Zen 2010 software (Zeiss). All antibodies used are listed in Supplementary Table…

### Immunofluorescence on paraffin sections

Neonatal hearts were fixed in 4% PFA for 24 hours at 4°C, washed three times with PBS, dehydrated through increasing concentration of ethanol, cleared with xylene, and finally embedded in paraffin. Hearts were then sectioned at 10μm. Hearts sectioned were dewaxed and rehydrated followed by antigen retrieval in preheated Sodium Citrate solution (pH= 6). The slides were then cooled down on ice and permeabilized using 0.5% PBT for 15 minutes at room temperature with agitation. Slides were washed with 1X PBS and incubated with a blocking solution (5% Donkey serum, 0.3% Tween- 20, 20mM MgCl2 in 1X PBS) for 1 hour at room temperature. Primary antibodies, diluted in above mentioned blocking solution, were incubate O/N at 4°C. The following day, slides were washed in 1X PBS and incubated with corresponding fluorescent-dye-conjugated secondary antibodies (5% BSA in 1X PBS) for 1 hour at room temperature. Next, slides were washed in 1X PBS, stained with DAPI for 15 minutes at room temperature, washed again with 1X PBS and mounted in Fluromount G (Southerm Biotech; Cat: 0100-01) prior to confocal imaging.

### Immunofluorescence on cryosections

Embryonic or neonatal hearts were fixed in 4% PFA for 1 hour at 4°C, washed in 1X PBS and dehydrate in 30% sucrose solution at 4°C O/N. The following day, hearts were embedded in OCT (Optical Cutting Temperature Compound, Fischer Health Care, Catalog: 4585), and stored at −80°C. 20μm cryosections were thawed at room temperature for 15 minutes followed by two consecutive washes in 0.5% PBT and incubated in blocking solution (5% Donkey serum, 0.5% PBT) for 1 hour. Primary antibodies were incubated at 4°C O/N. The following day, the slides were washed in 0.5% PBT, and incubated with corresponding fluorescent-dye-conjugated secondary antibody for 2-3 hours at room temperature followed by washes in 0.5% PBT. Cryosections were then mounted in Fluromount G prior to confocal imaging. For Cxcr4, staining was performed using tyramide signal amplification (TSA, Perkin Elmer, Catalog: NEL701A001KT).

#### In situ hybridization

*In-situ hybridization* for detection *Cxcl12* transcript was performed on paraffin sections as previously described (Koop et al. 1996)

### Neonatal LCA ligation

This procedure was adopted from Mahmoud et al. (Mahmoud et al. 2014). Briefly: P4 neonatal mice were gently wrapped in two layers of surgical gauze and placed on ice for 6 minutes to induce hypothermic circulatory arrest. Neonates were then orientated in a supine position on a cold pack covered with a plastic wrap. Before surgery, thoracic area was disinfected with iodine followed by 70% ethanol. Left anterolateral thoracotomy was made under dissecting microscope guidance. Dissection was carried through the pectoralis major and minor muscles, and the thoracic cavity was opened via the 4th intercostal space. The left coronary artery was identified and ligated using 8-0 prolene suture. The chest was then closed in layers with interrupted 7-0 prolene sutures. For AAVs related experiments, 50ml of rAAVs were injected into the thoracic cavity with an insulin syringe immediately after surgery. Neonates were then allowed to recover at 37°C warm plate and, when conscious, returned to its mother’s care. Due to mouse genetic inheritance, values for each parameter were compiled from both males and females from multiple litters.

### Production of AAVs

Gibson Assembly cloning method was used to insert a 2A peptide coding sequence immediately before the full-length of murine *Bmp2* (SinoBiological, Cat.: MG51115-G, NM_007553.2) and subsequently subcloned into pENN.AAV.cTNT.PI.eGFP.WPRE serotype 9 adenoviral vector (Addgene, Cat.: #105543). A modified protocol from Wakimoto et al. was used to produces AAVs (Wakimoto et al. 2016). HEK293T cells were transiently transfected with adenoviral helper plasmid pAdDeltaF6 (Addgene, Cat.:112867), trans-plasmid encoding AAV replicase and capsid gene pAAV2/9 (Addgene, Cat.:112865) and either pAAV-cTnT-GFP (vehicle) or pAAV-cTnT-GFP-p2A- Bmp2 vectors. 48-72hrs post-transfection, cells were harvested and centrifugated. AAVs particles were then purify from AAV-producing cells using AAVpro Purification Kit Maxi (Takara, Cat. #6666).

AAV titer from each sample was measured by Droplet Digital PCR (ddPCR).

### Embryonic ventricular explants

Ventricular explants were performed following modified version of the protocol from Large et al. (Large et al. 2020). The experiment was performed three times. E10.5 ventricles from each embryo were dissected in sterile 1X PBS and gently placed onto a cell culture insert (EMD Millipore, Cat. PI8P01250) coated with Matrigel (Corning, Cat. 354230). Two explants for each condition were located to the center of the insert. Excess of 1X PBS was removed and 200 μl of endothelial cell growth basal media (EBM-2, Cat. CC-3156) supplemented with 2% FBS (HyClone, Cat. SH3007003IR) were added to the space between the insert and the well. Explants were then covered with extra 100 μl of 2% FBS basal media. Ventricles were cultured in these conditions at 37 °C for 16 h. The day after, explants were washed with 1X PBS and recombinant rBMP2 (R&D Systems, Cat. 355-BM-010) or rVegfa (PeproTech, Cat. 450-32) diluted in 2% FBS basal media was added to the explants individually or in a 1:1 mixture at a final concentration of 20ug/ml. Explants cultured only with 2% FBS basal media represented vehicle controls. at 37 °C for approximately 4 days in basal media or experimental conditions basal media was replaced. Explants were incubated at 37 °C for 4 days replacing media every 24 hours. Finally, explants were washed with 1X PBS and fixed in 4% PFA for 10 min at 4 °C.

### Single-cell RNA sequencing protocol Processing of sequencing data

Raw Illumina reads for all datasets were demultiplexed and converted to FASTQ using bcl2fastq (Illumina). Reads were aligned to GRCm38 Ensembl release 81 as well tdTomato sequences and a gene count matrix was obtained using Cell Ranger v3.1.0 (10X Genomics).

### Processing of count data

The majority of scRNAseq data analysis was performed using R and Seurat v3 (Stuart et al. 2019). Cells were deemed low-quality and excluded from downstream analysis if they expressed less than 1000 genes or if more than 10% (E12, E17.5) of reads aligned to mitochondrial genes. A small number of cells were removed from the e17.5 tdTomato negative sample which were expressing tdTomato. For both datasets, non-endothelial subtypes (e.g., blood and immune cells, cardiomyocytes, smooth muscle, fibroblasts) as well as a small number of lymphatic cells were removed.

Normalization, variable feature selection, scaling, and dimensionality reduction using principal component analysis were performed using the standard Seurat v3 pipeline (Stuart et al., 2019). Technical variables genes per cell, reads per cell, cell cycle and mitochondrial read percentage were regressed out in the ScaleData function. Following this, construction of a shared nearest neighbour graph, cluster identification with the Louvain algorithm (Stuart et al., 2019), and Uniform Manifold Approximation and Projection (UMAP) dimensionality reduction (Becht et al., 2018) were performed using FindNeighbors (dims = 1:25), FindClusters (res=0.8), and RunUMAP (dims = 1:25) functions in Seurat.

A subset of ECs from E12 was used for further analysis (integration, trajectory and ligand-receptor interaction analysis): the endo EC clusters closest to CV ECs in the UMAP (E1 and E4), the SV cluster, and the CV clusters (CV1 and CV2). The VE cluster in E17.5 was excluded for the data integration and trajectory analysis.

### Differential expression testing

Differential gene expression testing in Fig. 5G was performed with the FindMarkers function in Seurat with the following parameters: logfc.threshold = 0.3, min.pct = 0.2 and filtered for p-value < 0.001.

### Datasets integration

After QC filtering, the samples were normalized, scaled and centred, 5,000 shared highly variable genes were identified using Seurat FindVariableFeatures and SelectIntegrationFeatures() functions. Integration anchors were identified based on these genes using canonical correlation analysis as implemented in the FindIntegrationAnchors () function with the following settings: reduction = ”cca”,dims = 1:5, k.anchor = 20. The data were then integrated using IntegrateData() and scaled again using ScaleData(). PCA and UMAP with 30 principal components were performed. A nearest-neighbour graph using the 30 dimensions of the PCA reduction was calculated using FindNeighbours(), followed by clustering using FindClusters() with a resolution of 0.5.

Same parameters have been used to perform integration all Endo ECs (Suppl. Figure 8B) and capillary cells (Suppl. Figure 8C) from E12 and E17.5.

### Trajectory analysis

We used the Bioconductor package Slingshot (Street et al. 2018), Dynverse (Saelens et al. 2019) within ASC Seurat web application (Pereira et al. 2021) to determine pseudotime lineage trajectories and identify top 50 genes DEG along the trajectory. CytoTRACE (Cellular (Cyto) Trajectory Reconstruction Analysis using gene Counts and Expression) (Gulati et al. 2020) to predict the differentiation state of the cells in the selected E12 and E17.5 clusters with default parameters.

### Ligand-receptor interaction analysis

For ligand-receptor analysis, we used CellChat (Jin et al. 2021) (https://github.com/sqjin/CellChat) v1.1 using the SecretedSignaling subset of the mouse CellChatDB, with default parameters.

Custom code was generated using R to analyze the data and to generate plots.

### Quantification and statistical analysis

Quantification of tdTomato expressing cells in ECs (tdTomato^+^-Erg^+^ cells) was performed from different z-stack in whole-mount images or heart sections at different time points. Total number of Erg+ cells was automatically calculated using Image-based Tool for Counting Nuclei (ITCN) plugin in FIJI whereas tdTomato+ cells were manually counted using the CellCounter plugin. In Fig. 1F and 6H, endo derived CVs contribution is represented as ratio between total number of tdTomato*+* cells and Vegr2 or IB4 vessel area measured using the AnalyzeParticle plugin in FIJI.

Colocalizaton images were produced using the “analyze particles” plugin in FIJI on masks of GFP+ nuclei and ERG+ nuclei chosen from a default threshold with parameters of circularity >.2 and area > 75 pixels. The two masks were combined using the AND function in the RIO manager and the resulting pixels were overlaid on the TD-Tomato expressing channel.

Statistical analysis and graphs were generated using Prism 8 (GraphPad). All statistical tests were performed using two-sided, unpaired Student’s t-tests except. *P < 0.05, **P < 0.01, ***P < 0.001 and **** P ≤ 0.0001. Sample size was chosen empirically according to previous experience in the calculation of experimental variability. No statistical method was used to predetermine sample size. All experiments were carried out with at least three biological replicates. Numbers of animals used, and statistical significance are described in the corresponding figure legends.

## Contributions

G.D and K.R-H- conceptualized the study and wrote the article. G.D., R.P, and J.A.N. performed experiments. G.D. and A.V performed computational analysis. G.D., K.R-H., K.E.Q, and K.M.C. designed *Cxcr4-Tango* mouse and provided experimental resources. D.O.C advised on and generated *Cxcr4-Tango* transgenic line. B.S. provided advice on experiments and manuscript.

P.E.R.C. assisted with the neonatal injured experiments. All authors reviewed the manuscript.

## Acknowledgements

K.R.-H. is supported by the NIH/NHLBL (R01-HL128503). G.D. held an EMBO postdoctoral fellowship (ALTF 1542-2016) and is supported by NIH (T32HL120824). R.P. is supported by an AHA graduate fellowship. P.E.R.C is supported by the NIGMS of the National Institutes of Health (NIH T32GM007276) and NSF-GRFP (DGE-1656518). A.V. is supported by Biomedical Research Centre award and NIH. We thank Ralf Adams for sharing the *BmxCreER* mouse line. We thank José Luis de la Pompa for providing *ISH* probes. We thank Rahul Sinha for experimental assistance with FACS sorting. We thank Jiyeon Ban and Chris Cook for mouse breeding assistance. We thank all members of the Red-Horse lab for technical and intellectual support. We thank members of the Stanford Genome Sequencing Services Center which is supported by NIH Grant No. 1S10OD020141.

---

**Figure S1.**
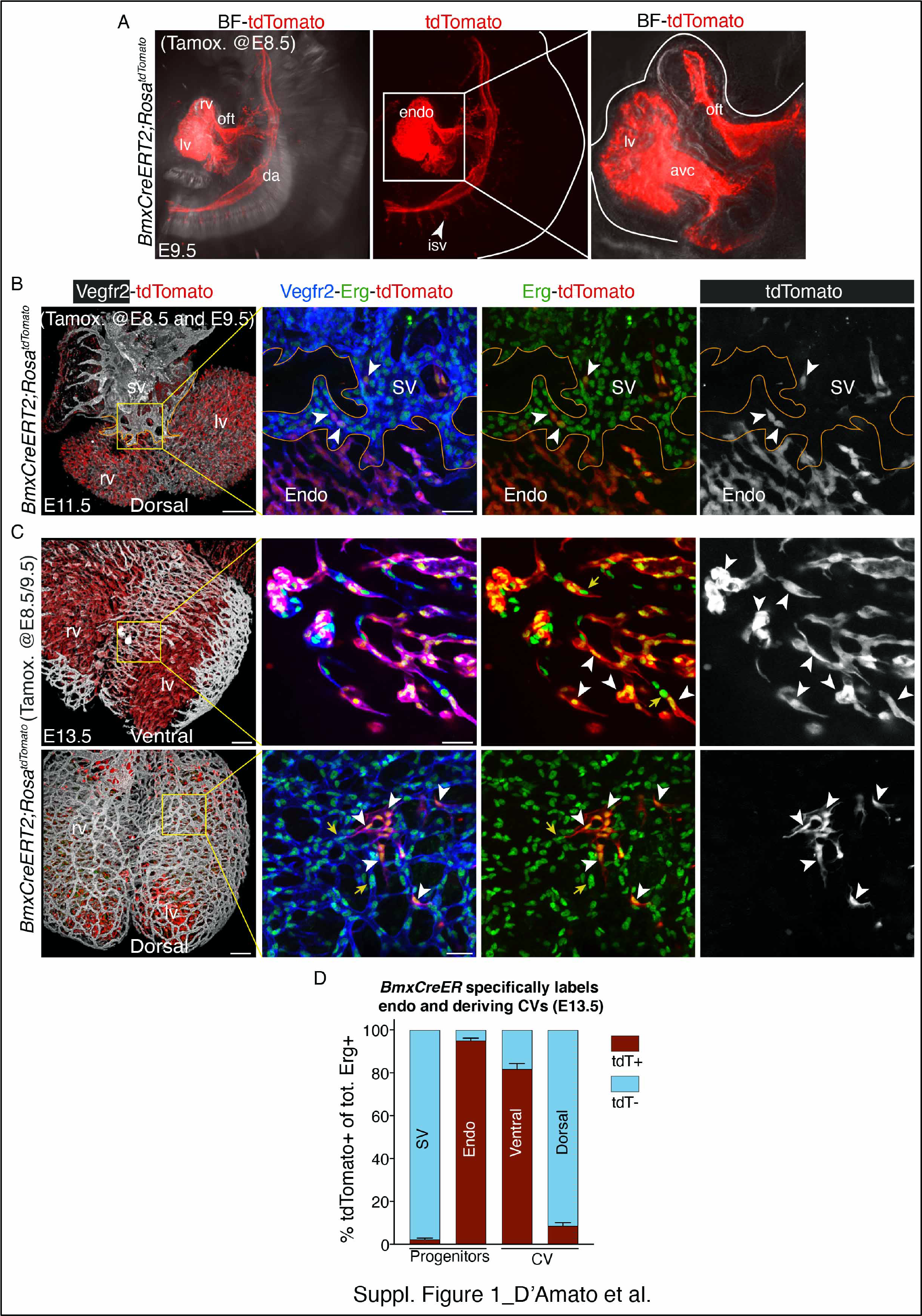
**A)** E9.5 light-sheet whole-mount images of *BmxCreERT2;Rosa^tdTomato^* tamoxifen- induced at E8.5 show expression of tdTomato (red) in endo and dorsal aorta. tdTomato expressing cells are also found in intersomitic vessels (isv, arrowhead). Magnification of E9.5 heart is shown in boxed region (avc=atrioventricular canal, oft=outflow tract; lv= left ventricle; rv= right ventricle). **B)** Whole-mount images of lineage traced E11.5 *BmxCreERT2;Rosa^tdTomato^* heart (dorsal view) immunostained Vegfr2 and Erg. Magnification (yellow box) show minimal contribution of endo-derived CVs (tdTomato^+^ cells, arrowheads) in SV and SV derived ECs (solid orange line). **C)** Ventral (upper panels) and dorsal (lower panels) view of E13.5 endo lineage traced heart. Boxed regions show the contribution of endo-derived cells to ventral and dorsal vascular plexus (arrowheards). **D)** Quantification in percentage of the total number of tdTomato^+^/Erg^+^ cells (red bar) from E13.5 *BmxCreERT2;Rosa^tdTomato^* heart sections (*n* = 4 hearts) in progenitors (endo and SV) and CV plexus (ventral and dorsal). Scale bar=200 μM in B and C (full view). Scale bar=50 μM in B and C (boxed regions).

**Figure S2.**
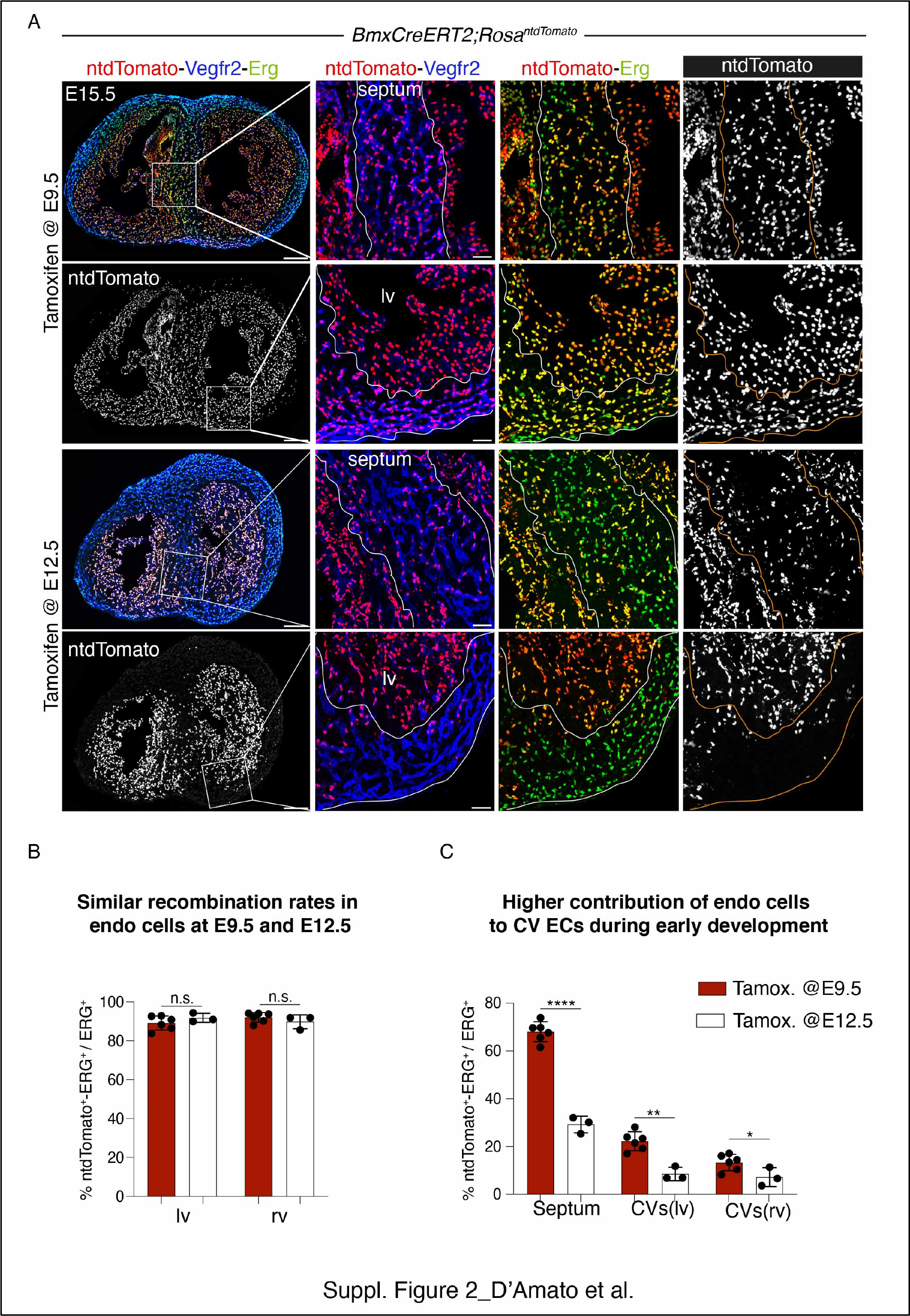
**A)** E15.5 Heart sections from of *BmxCreERT2;Rosa^ntdTomato^* immunostained for Vegfr2 (blue) and Erg (green). Tamoxifen was induced at E9.5 or E12.5. Boxed regions are magnification of septum and left ventricle (lv) respectively. In E12.5-induced hearts, *ntdTomato* (red) expression is greatly reduced in CVs of septum and ventral side of lv. **B and C)** Quantification in percentage of ntdTomato^+^ expressing cells in endo and ECs (Erg^+^) from E15.5 embryos injected at E9.5 (red bars, *n=6 hearts*) and E12.5 (white bars, *n=3 hearts*). Scale bar=200 μM in A and 50 μM in the boxed regions. Data are mean ± s.d. * *P* ≤ 0.5, \*\**P* ≤ 0.01, ****, P ≤ 0.0001, by Student’s *t*-test.

**Figure S3.**
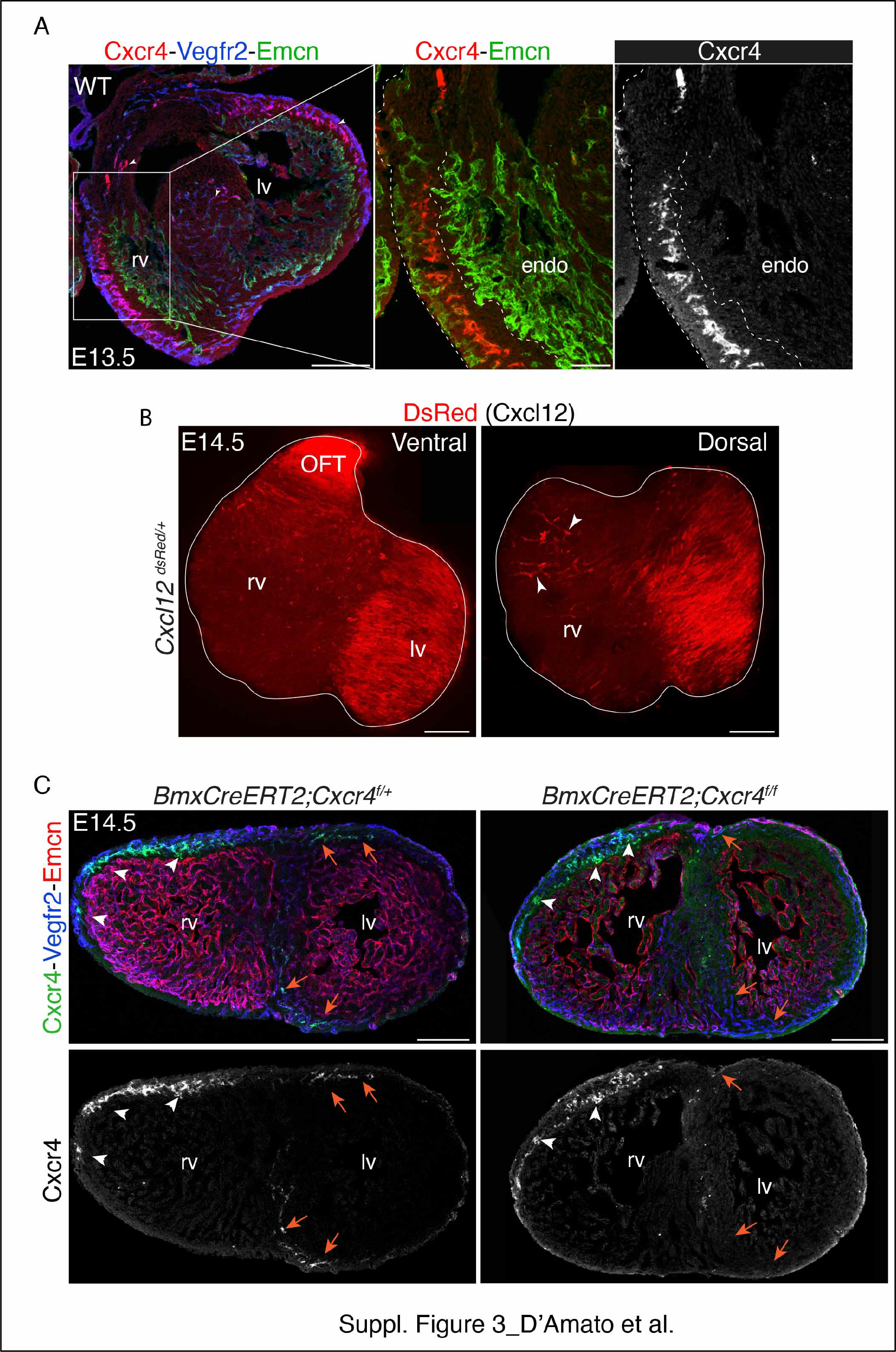
**A)** Confocal images of E13.5 WT heart section immunostianed for Cxcr4 (red), Vegfr2 (blue) and endocardial marker endomucin (Emcn, green). Cxcr4 is restricted in intramyocardial vessels and in not expressed in endo (boxed region). **B)** Ventral and dorsal view of E14.5 *Cxcl12^dsRed^* heart show *dsRed,* as a surrogate for *Cxcl12*, expressed in the left ventricle (lv) and outflow tract (oft). Arrowheads indicate expression in pre-artery CVs. **C)** E14.5 heart sections from *BmxCreERT2;Cxcr4^f/+^* (control) and *BmxCreERT2;Cxcr4^f/ f^* (mutant) immunostained for Cxcr4 (green), Vegfr2 (blue) and Endomucin (Emcn, red). White arrowheads show SV derived ECs expressing Cxcr4. Orange arrows indicate Cxcr4 expression in endo derived CVs that is lost in mutant heart. Scale bar=200 μM in A, B and C (full view). Scale bar=50 μM in A (boxed regions).

**Figure S4.**
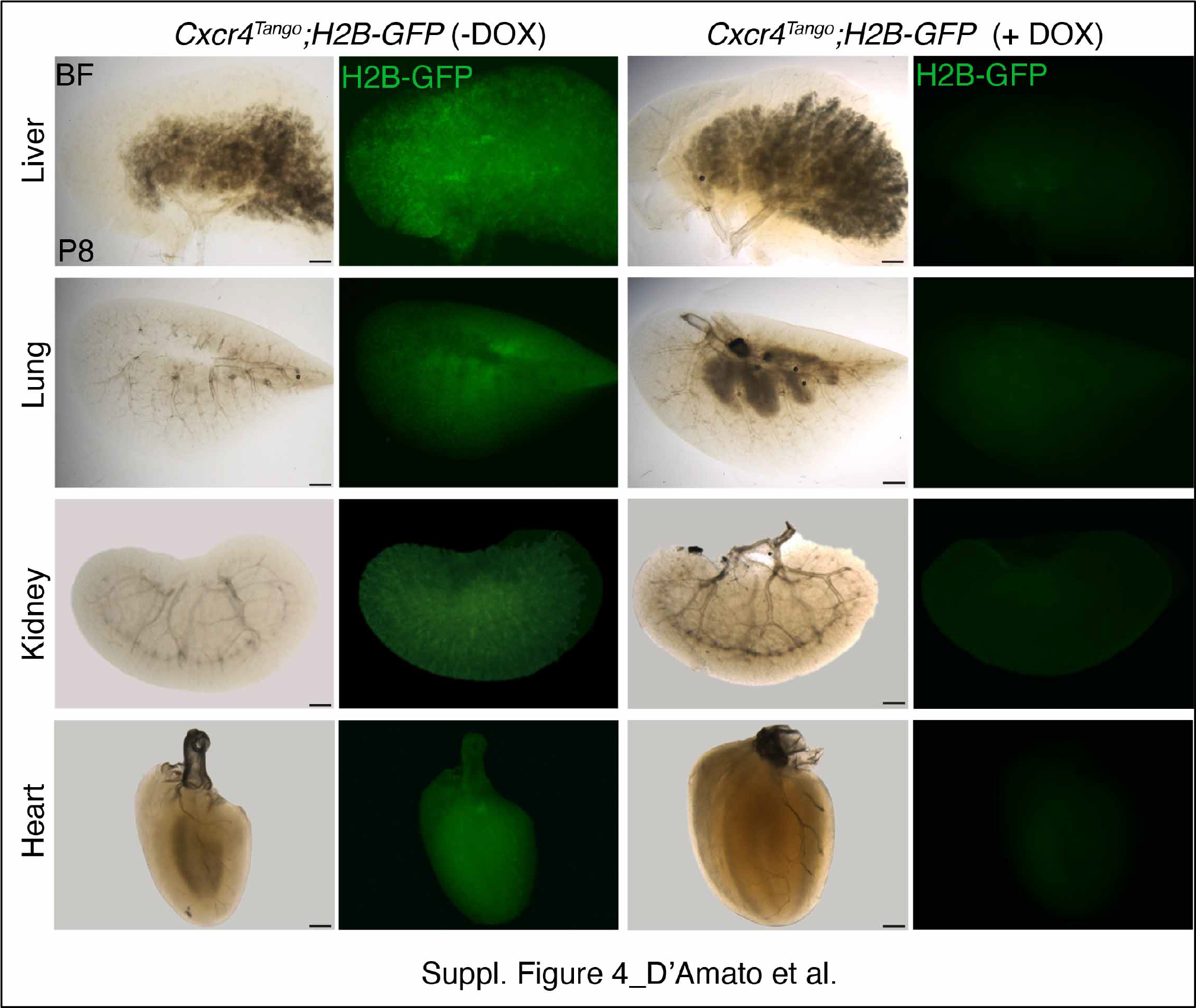
**A)** Stereoscopic images of P8 liver, lung, kidney and heart from *Cxcr4^tango^;H2B-GFP* animals with (+ Dox) and without (- Dox) doxycycline treatment. Cxcr4 activity, as surrogate of H2B-GFP expression (green), is detected only in non-treated animals. Organs are arrange based on their fluorescent intensity (top panels=strongest fluorescent intensity; bottom panels=weaker fluorescent intensity). Scale bar=200 μM.

**Figure S5.**
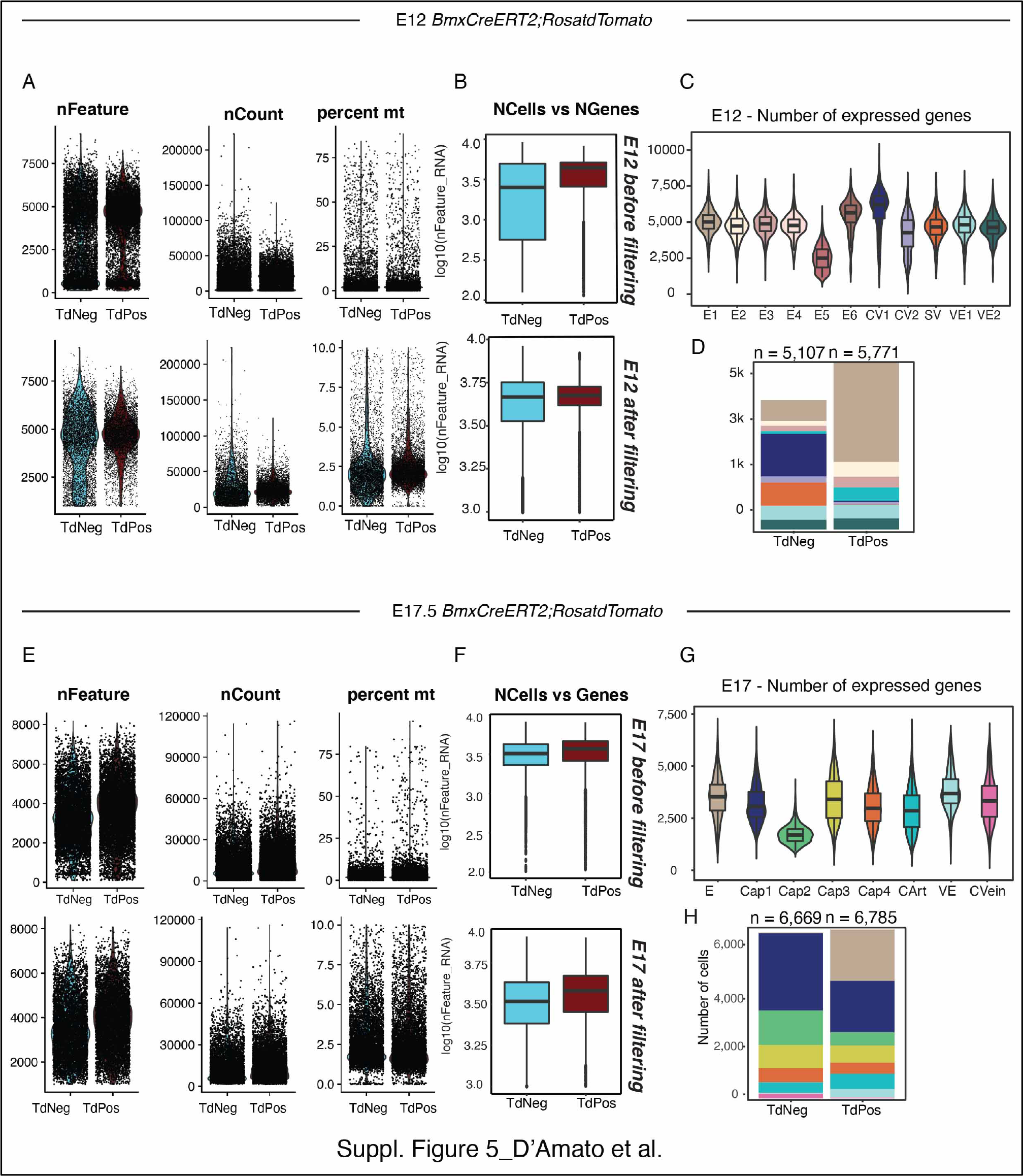
**A)** Violin plots of quality control metrics used to filter cells in E12 dataset before analysis (top panels). Cells were filtered for number of total aligned reads and percent mitochondrial reads (nFeature > 1000 and <10% mitochondrial reads [pct.mt]) (bottom panels). **B)** Boxplots showing the distribution of genes detected in tdTomato+ and tdTomato- E12 cells before (top) and after (bottom) filtering. **C)** Violin plots showing the average number of expressed genes for every cluster in the E12 dataset. **D)** Barplot depicting composition of E12 clusters in each sample (tdTomato+ and tdTomato-) by actual number of cells. **E)** Violin plots of quality control metrics used to filter cells in E17 dataset before analysis (top panels). Cells were filtered for number of total aligned reads and percent mitochondrial reads (nFeature > 1000 and <10% mitochondrial reads [pct.mt]) (bottom panels). **F)** Boxplots showing the distribution of genes detected in tdTomato+ and tdTomato- E17 cells before (top) and after (bottom) filtering. **G)** Violin plots showing the average number of expressed genes for every cluster in the E17 dataset. **H)** Barplots depicting composition of E17 clusters in each sample (tdTomato+ and tdTomato-) by actual number of cells.

**Figure S6.**
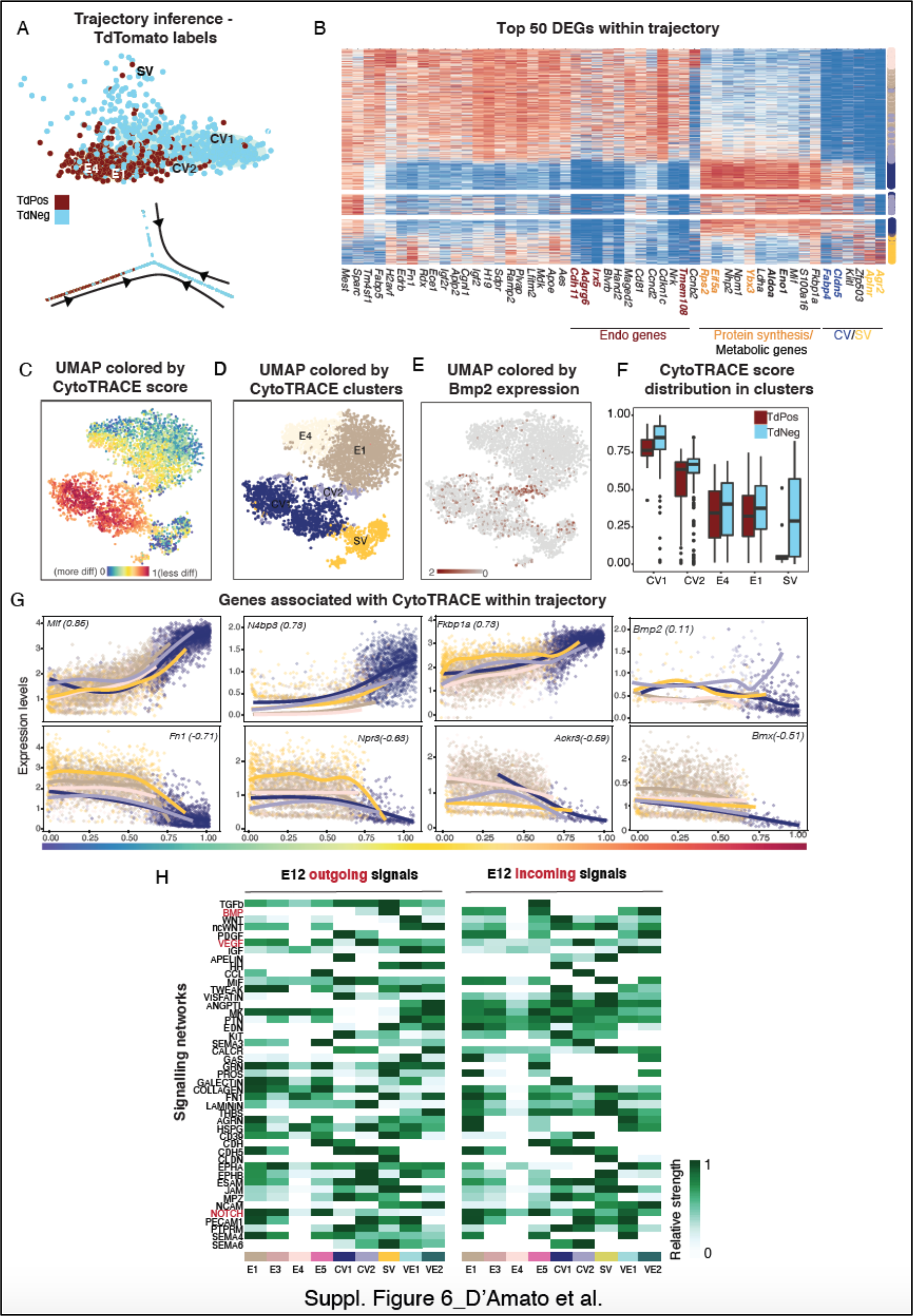
**A)** Visualization of the trajectory inferred using slingshot on a dimensionality reduction of the cells (top panel) and as a 2D graph structure (bottom panel) colored by tdTomato identity. **B)** Heatmap showing the expression of the 50 most significant DEGs within the trajectory, sorted by their position in the inferred trajectory. **C)** The Uniform Manifold Approximation and Projection (UMAP) obtained by applying CytoTRACE to the E1-E4-CV1-CV2-SV clusters, colored by CytoTRACE score, (**D**) cluster identity and (**E**) Bmp2 expression. **F)** Boxplots indicating the Cytotrace score distribution in Tdtomato+ and TdTomato- cells in each cluster. **G)** Scatter plots showing correlation between expression of genes (y-axis) associated with stemness and differentiation and their CytoTRACE score (x-axis). Each cell is colored by cluster identity. Gene names and Pearson correlations score are indicated. **H)** Major ligand/receptor (L/R) relationships in the E12 dataset displayed as a heatmap showing outgoing ligands (left) and incoming receptivity (right). Overrepresented signalling pathways are along left y-axis, and cell identity is along bottom x-axis, with colors corresponding to cell identity.

**Figure S7.**
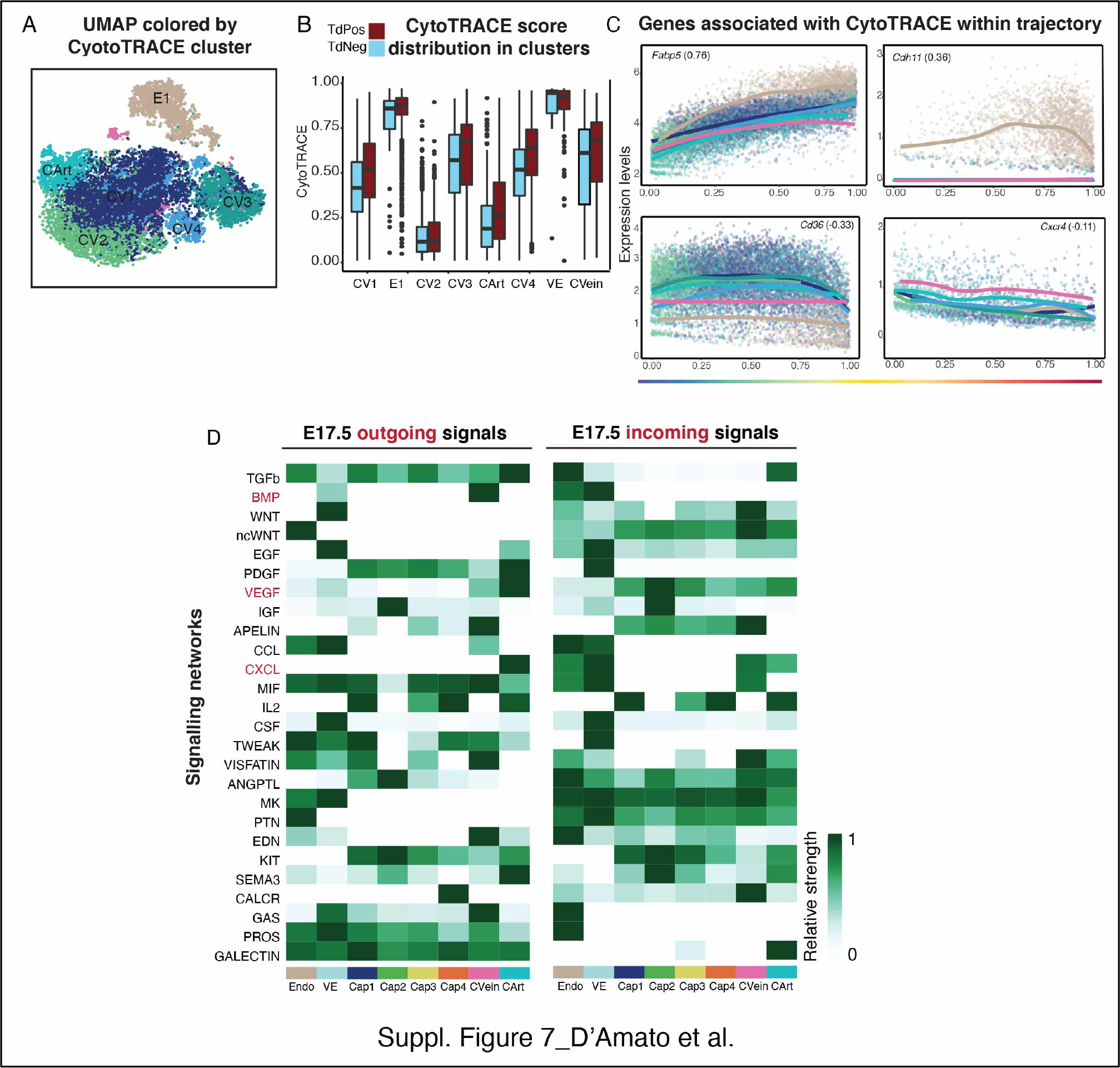
**A)** The Uniform Manifold Approximation and Projection (UMAP) obtained by applying CytoTRACE, colored by cluster identity. **B)** Boxplots indicating the Cytotrace score distribution in Tdtomato+ and TdTomato- cells in each cluster. **C)** Scatter plots showing correlation between expression of genes (y-axis) associated with stemness and differentiation and their CytoTRACE score (x-axis). Each cell is colored by cluster identity. Gene names and Pearson correlations score are indicated. **D)** Major ligand/receptor (L/R) relationships in the E17.5 dataset displayed as a heatmap showing outgoing ligands (left) and incoming receptivity (right). Overrepresented signalling pathways are along left y-axis, and cell identity is along bottom x-axis, with colors corresponding to cell identity.

**Figure S8.**
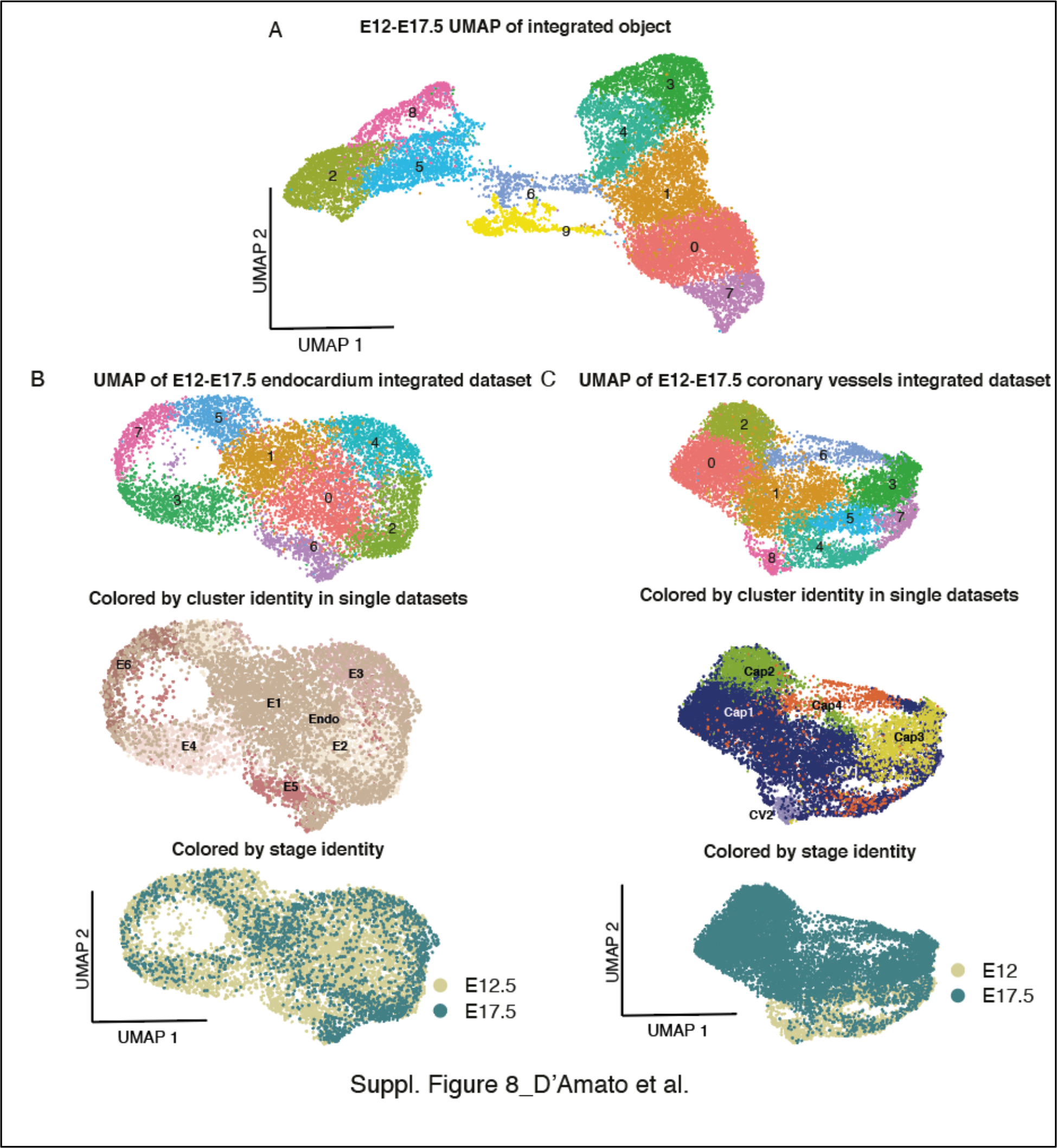
**A)** The Uniform Manifold Approximation and Projection (UMAP) of the 9 identified main clusters in the integrated E12-E17 dataset. **B)** UMAPs showing integration of Endo ECs across the two datasets, colored by cluster (top), cluster identity in single datasets (middle) and dataset (bottom). **C)** UMAPs showing integration of coronary vessel clusters across the two datasets, colored by cluster (top), cluster identity in single datasets (middle) and stage identity dataset (bottom).

**Figure S9.**
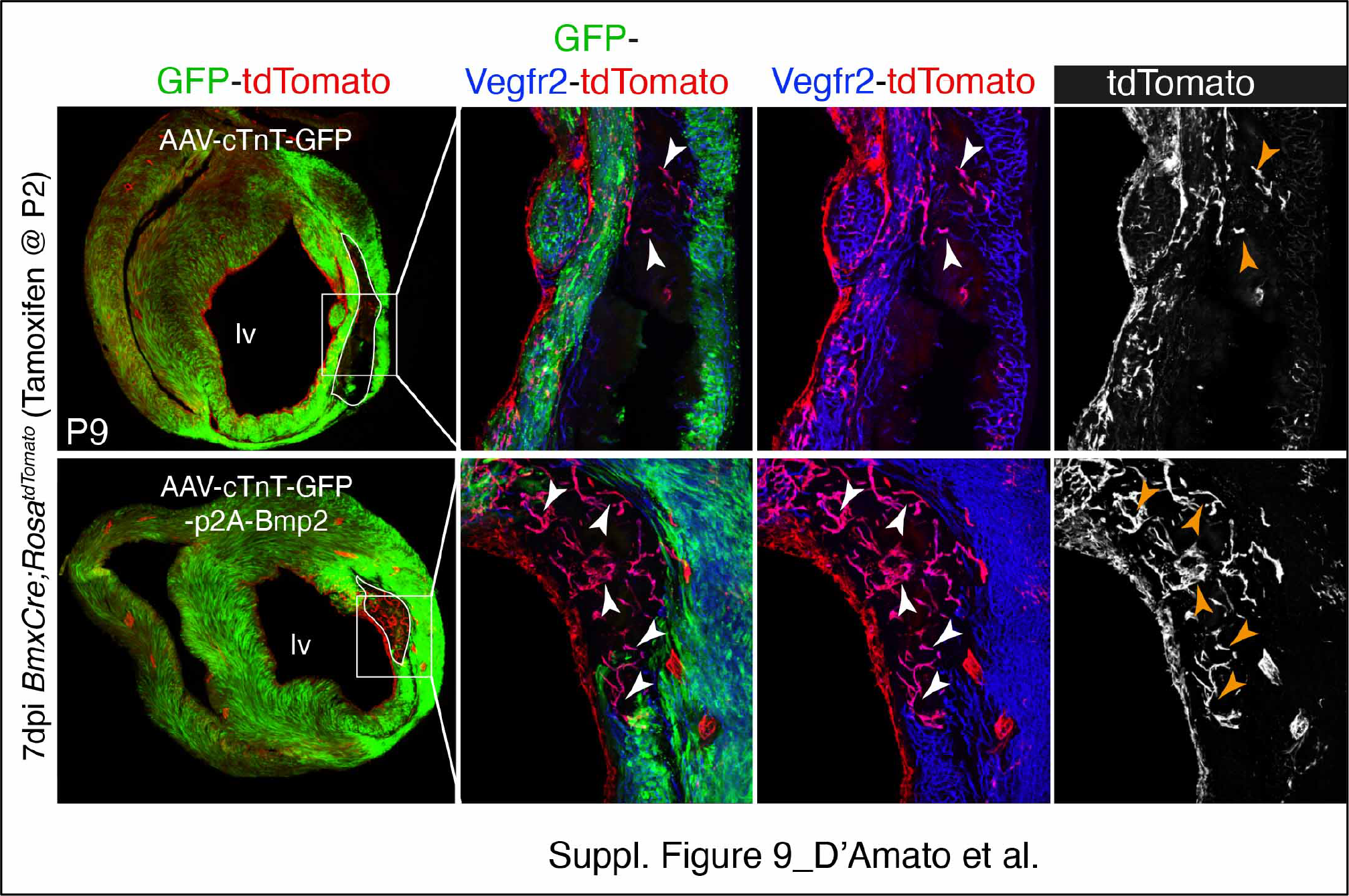
Immunostaining of *BmxCreERT2;Rosa^tdTomato^* P9 heart sections one week after MI AAVs administration. GFP (green) is expressed in cardiomyocytes of AAVs transduced hearts. Boxed regions show higher number of endo-derived CVs (tdTomato+ in ECs, arrowheads) in the scar area of Bmp2-trasduced heart (lower panel) compared with control heart (upper panel). Scale bar=200 μM (full view) and 50 μM in boxed regions. Scale bar=50 μM in A (boxed regions).

