## Supplementary figures and images for "Endocardium-to-coronary artery differentiation during heart development and regeneration involves sequential roles of Bmp2 and Cxcl12/Cxcr4"

### Supplementary Figure 1

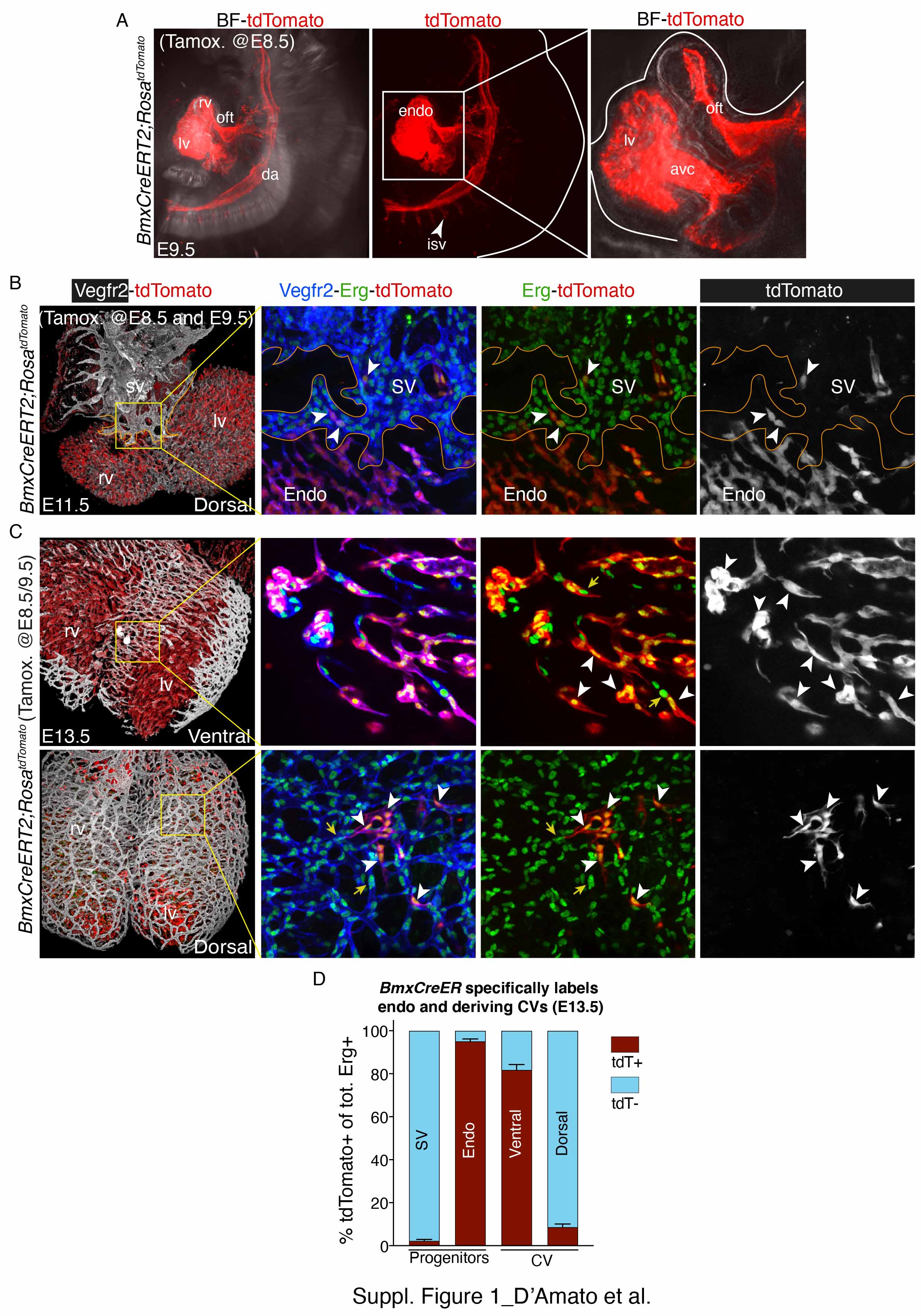

### Supplementary Figure 2

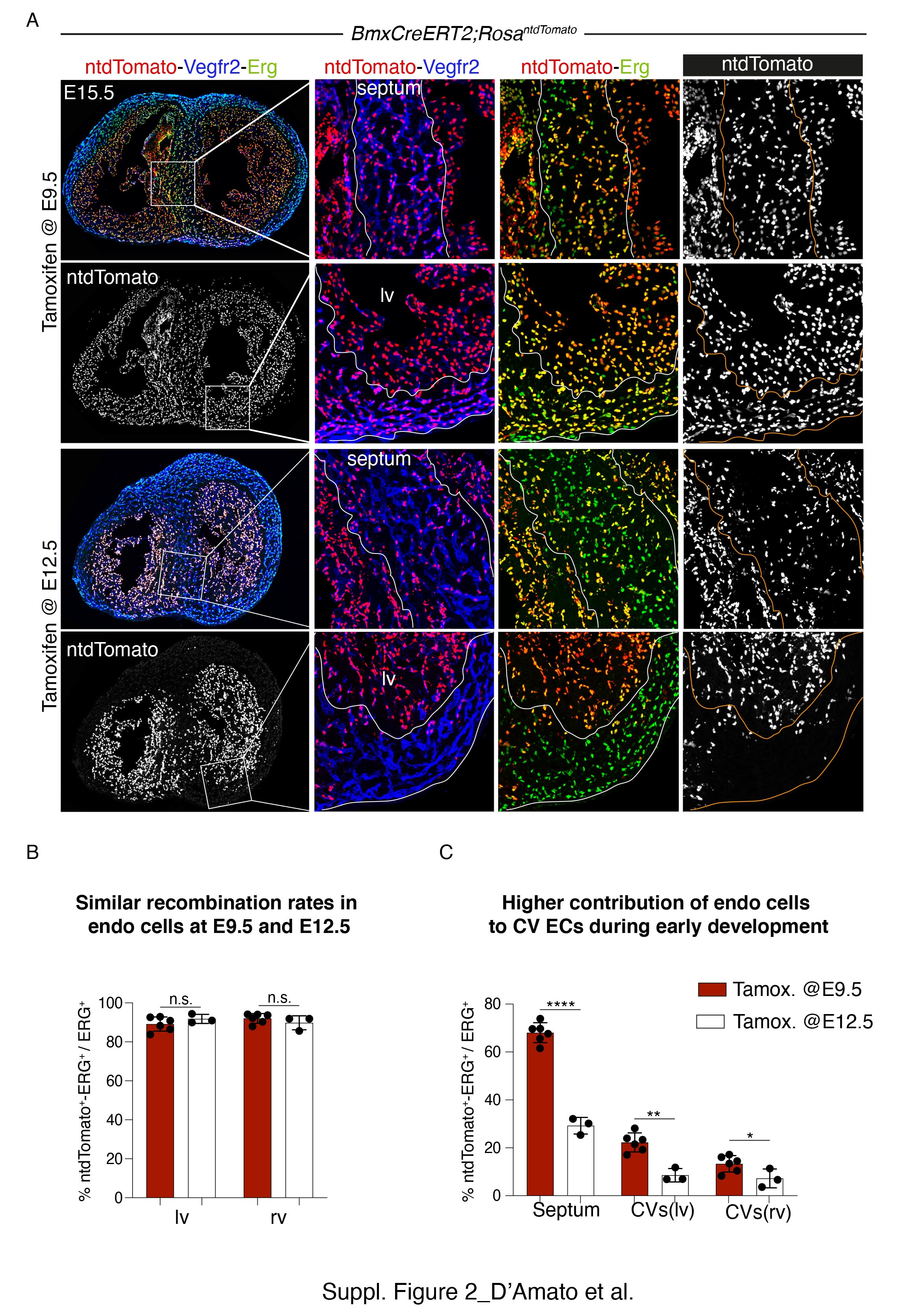

### Supplementary Figure 3

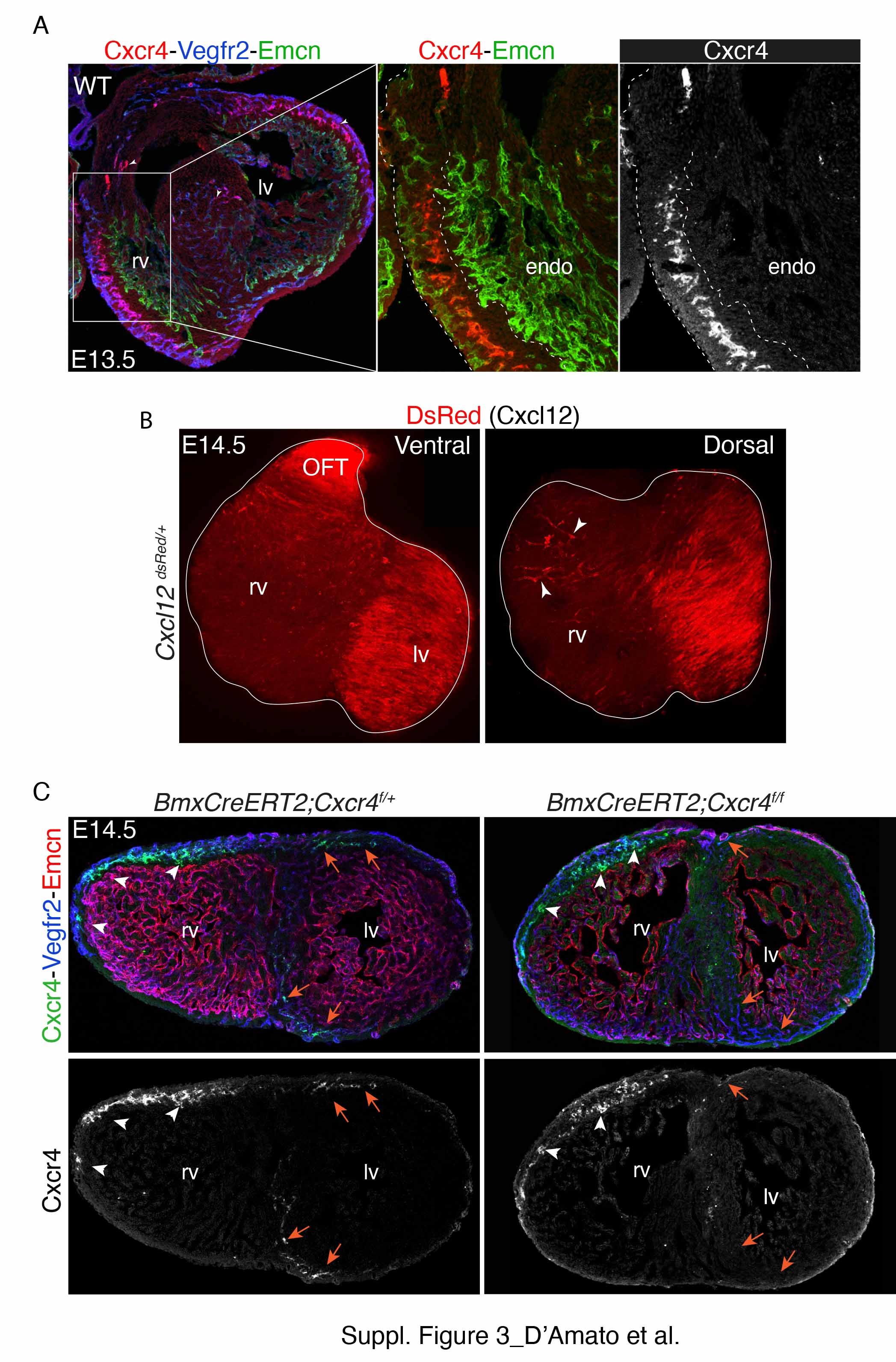

### Supplementary Figure 4

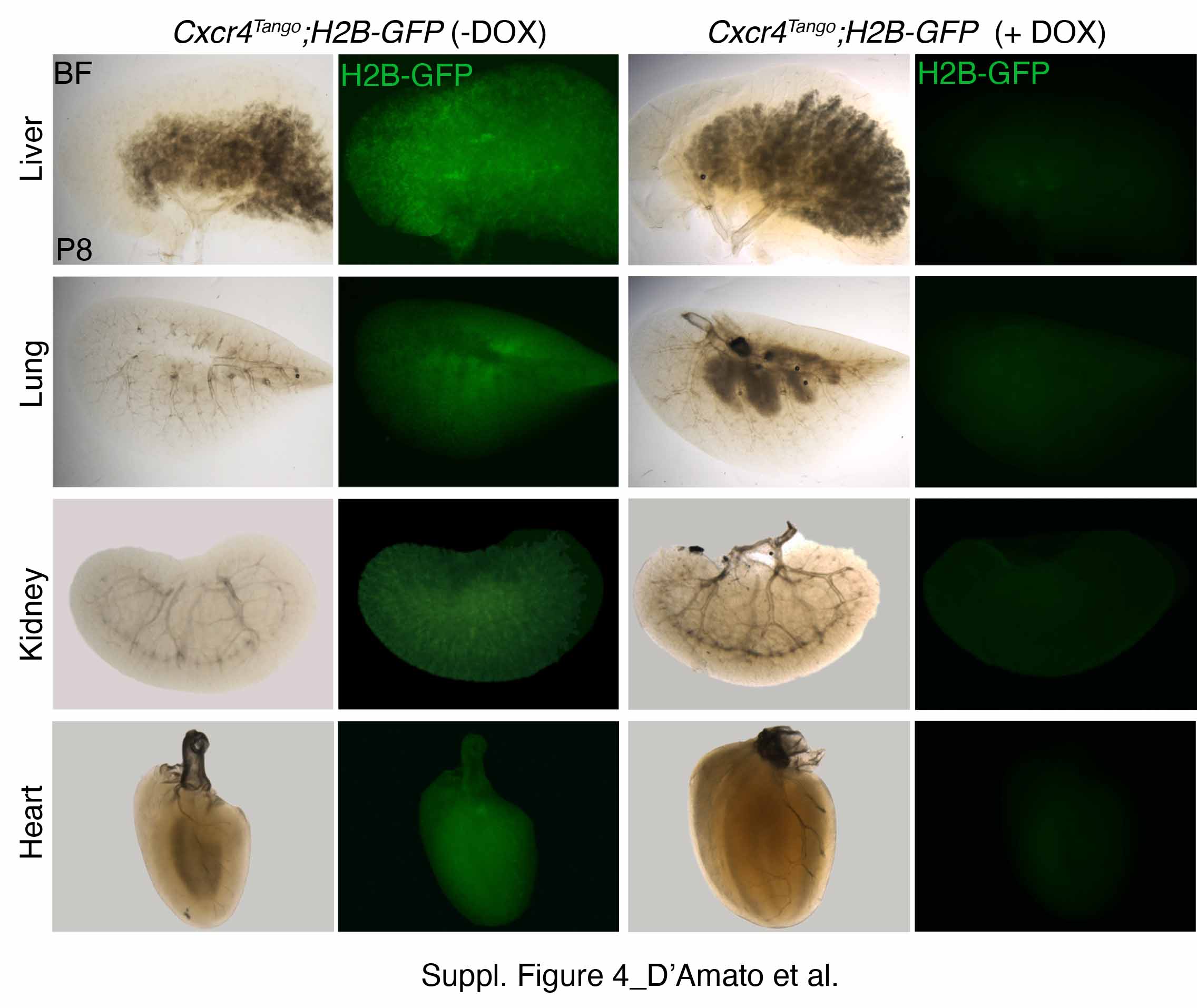

### Supplementary Figure 5

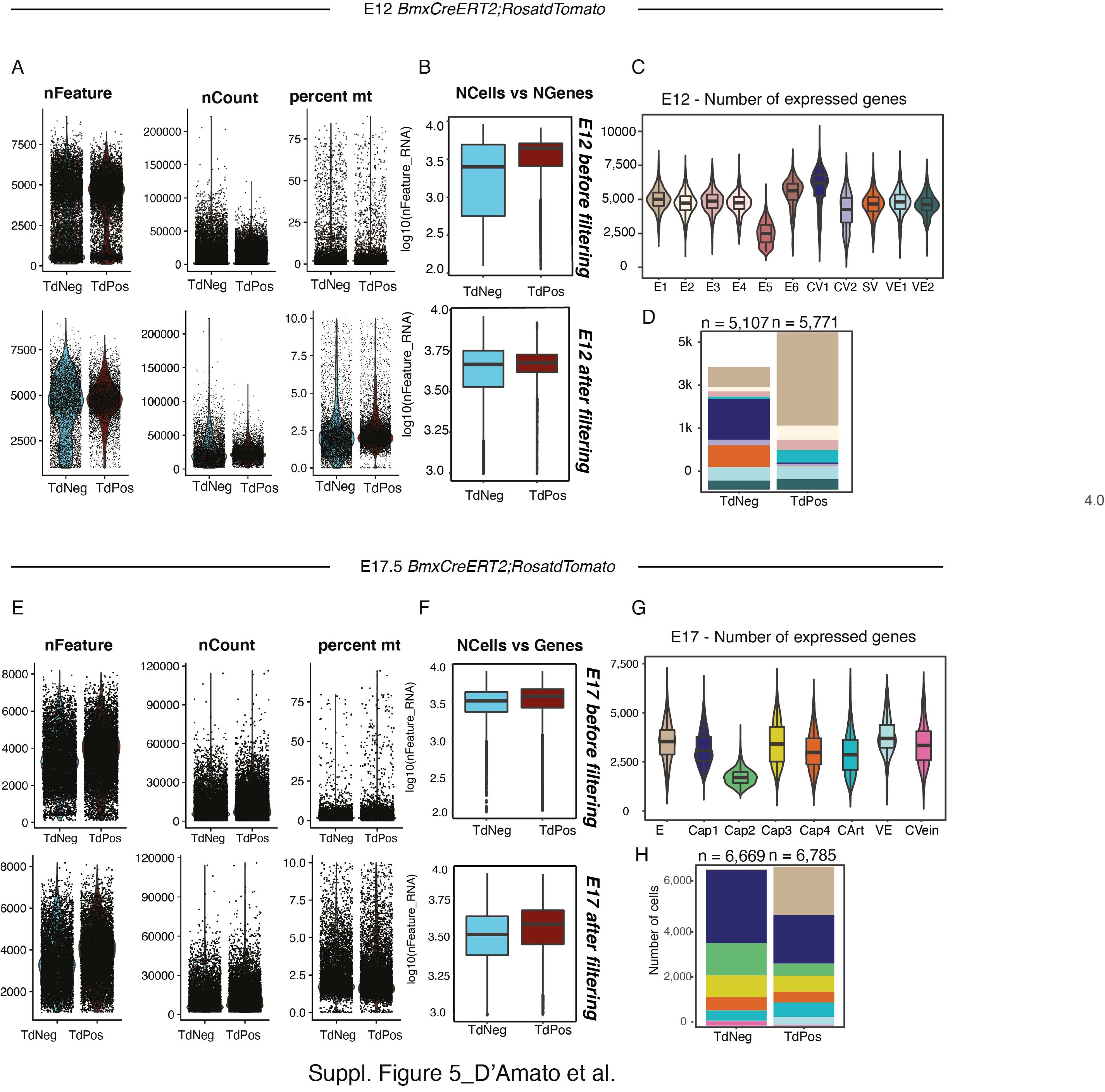

### Supplementary Figure 6

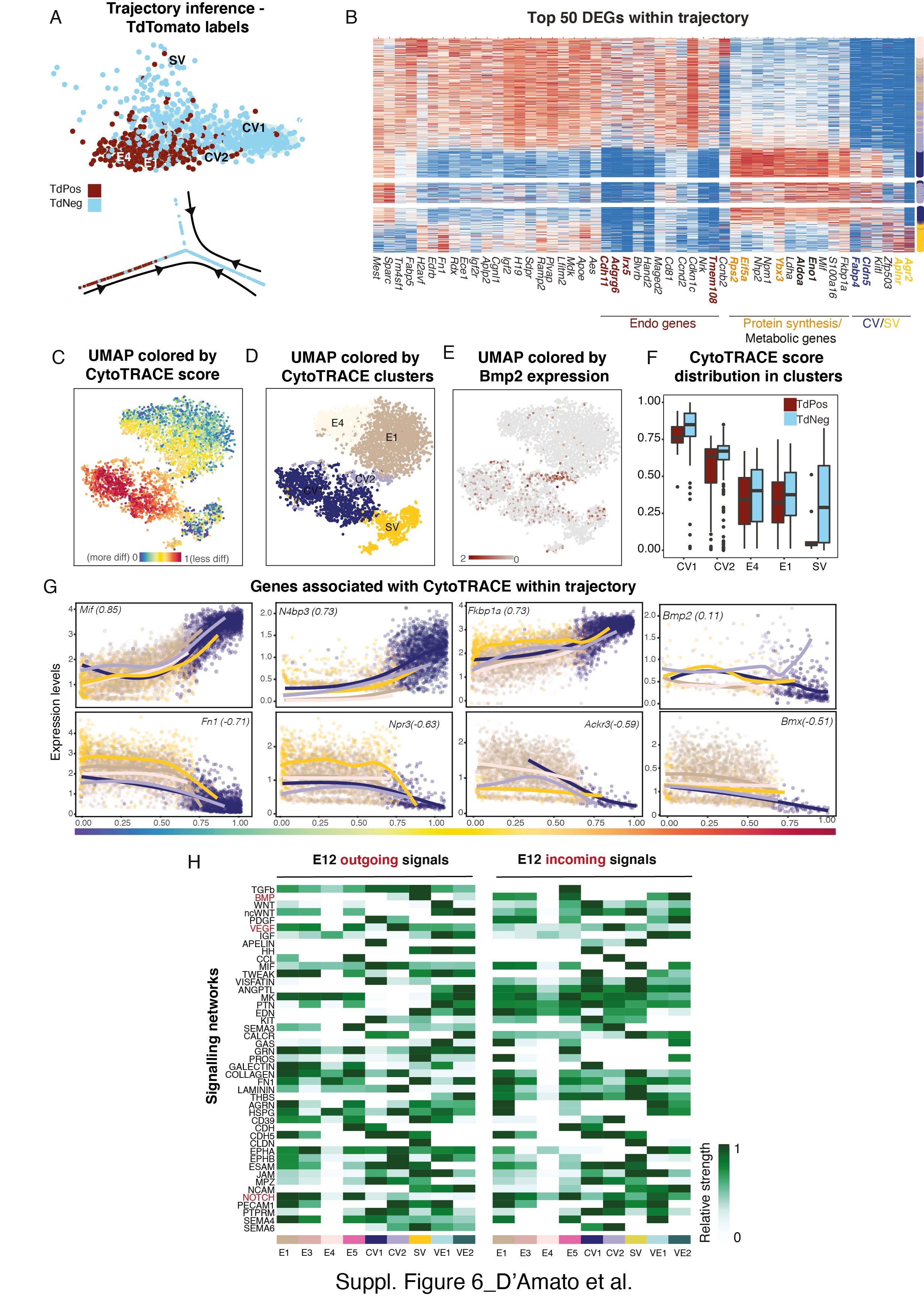

### Supplementary Figure 7

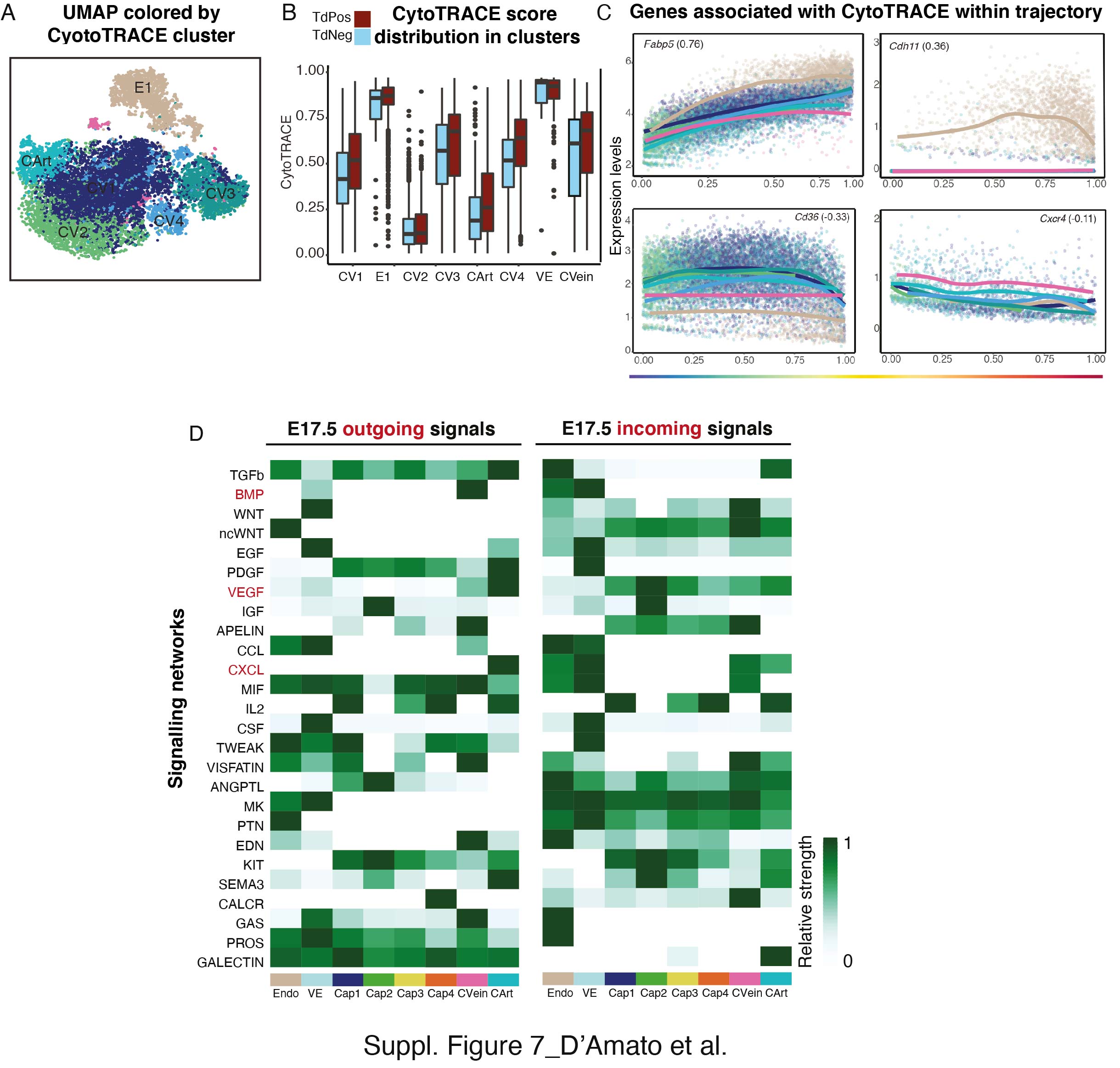

### Supplementary Figure 8

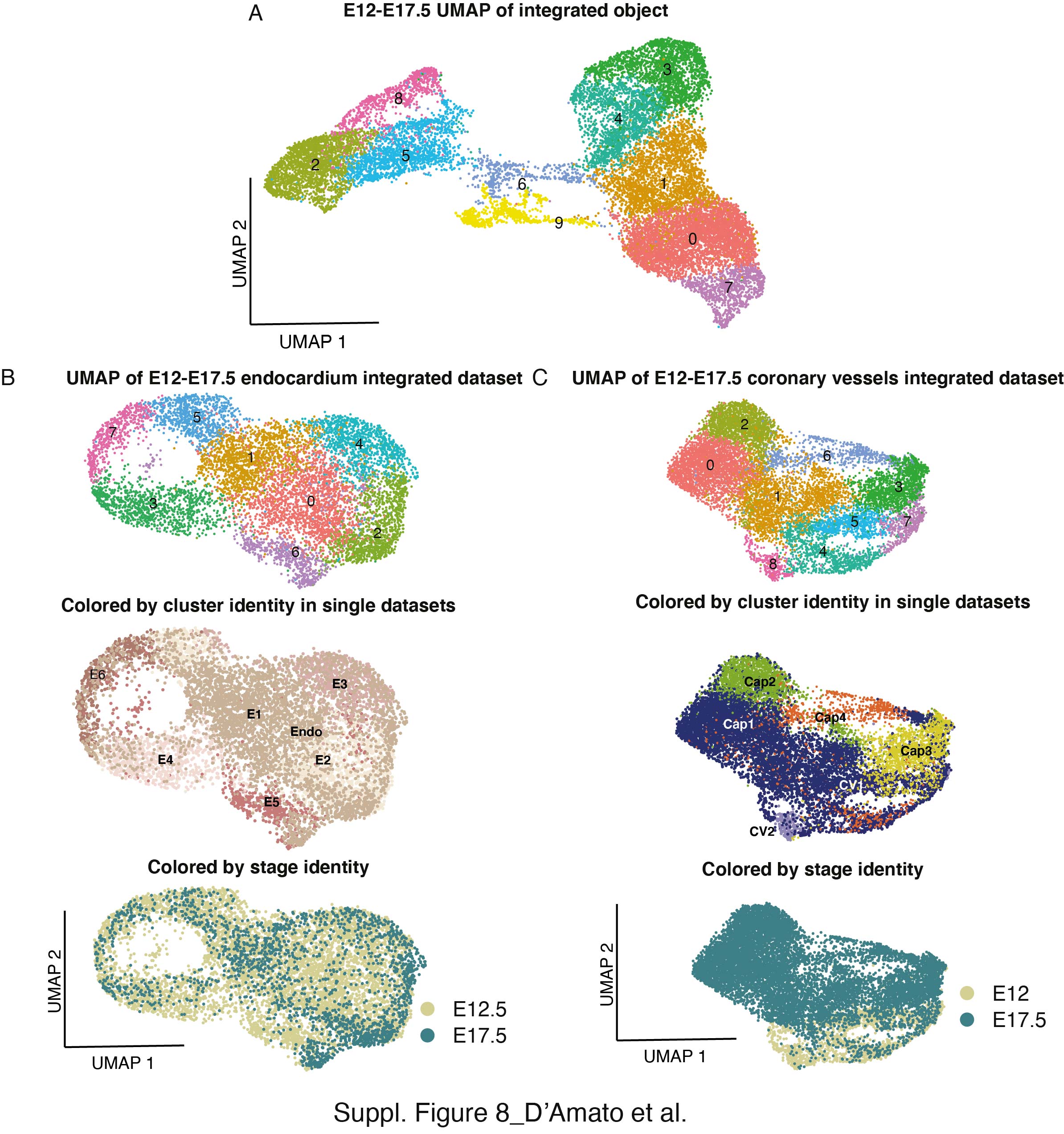

### Supplementary Figure 9

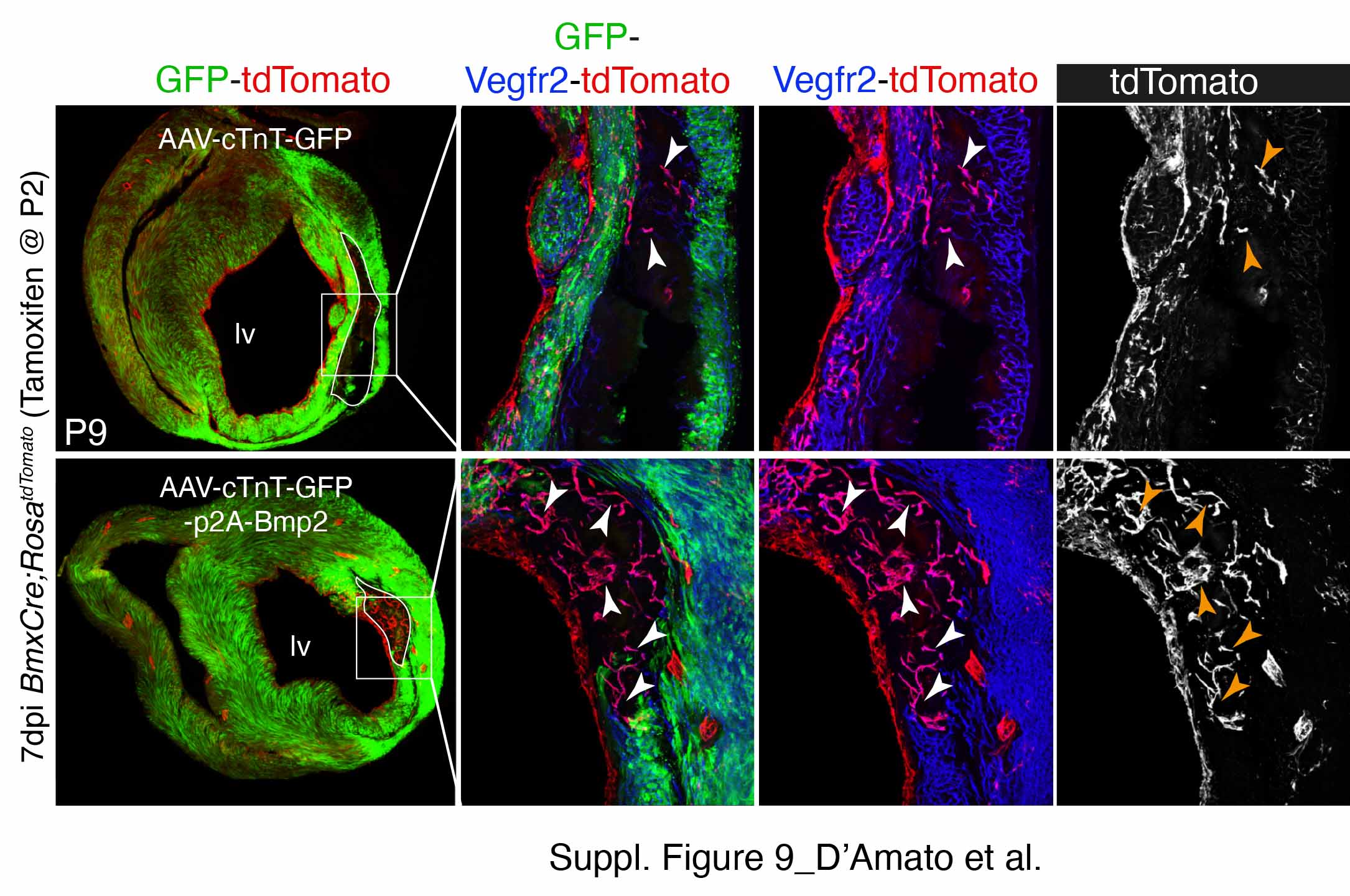
